# SOX9-mediated G1 elongation confers reserve stem cell-associated injury resistance in human intestinal stem cells

**DOI:** 10.64898/2026.08.21.745750

**Authors:** Joseph Burclaff, Keith Breau, Lauren T. Chi, Whitney DeLoach, Ebenezer A. Amare, Lauryn Cooper, Vanessa Walcott, Caroline Hinesley, Michelle Dixit, Kaiwen Chen, Mickael Meyer, Caden Sweet, Davis Walker, R. Jarrett Bliton, Cynthia Y. Tang, Scott T. Magness

**Author notes:** **Address Correspondence To:** Scott T. Magness, PhD, Department of Biomedical Engineering, University of North Carolina at Chapel Hill, 111 Mason Farm Road, Chapel Hill, NC, USA., Joseph Burclaff, PhD, Department of Molecular Biomedical Sciences, College of Veterinary Medicine, North Carolina State University, 1060 William Moore Drive, Raleigh, NC, USA.

## Abstract

**Background & Aims:** Dynamic cell cycle control is critical for intestinal crypt maintenance and injury responses, yet genetic regulators driving these changes remain poorly defined. As reserve intestinal stem cells (rISCs) are often considered to be slowly-cycling and can resist replication-dependent injury, factors that restrain proliferation may confer cytoprotection. Here, we define SOX9 as a regulator of intestinal stem cell (ISC) cycling and injury resistance.

**Methods:** Primary human ISCs were engineered to tune SOX9 levels, visualize cell cycle state, and manipulate cell cycle regulators. Using this system, we tested how SOX9 dosage impacts stemness, differentiation, proliferative recovery after SOX9 washout, and survival after 5-FU-mediated injury. Transcriptional analyses identified candidate links between SOX9 levels and cell cycle control, which were functionally tested using inducible INK4A (CDKN2A) and Cyclin D2 (CCND2) ISC lines.

**Results:** SOX9 induction lengthens the cell cycle in a dose-dependent manner largely by elongating G1 phase through the INK4A-Rb pathway. The effects of high SOX9 levels repressing proliferation and stem cell activity are reversible. SOX9 induction protects against 5-FU toxicity. This protection is mimicked by INK4A overexpression or pharmacological G1 phase arrest and repressed by CCND2 induction.

**Conclusions:** These findings identify SOX9-mediated G1 elongation as a reversible cytoprotective program that confers key functional properties associated with rISCs: proliferative restraint, retained stem cell potential, and resistance to replication-dependent injury. This positions G1 length as a potential determinant of which crypt cells survive injury to act as reserve stem cells.

## Introduction

The mammalian small intestine is one of the most proliferative tissues in the body, with near complete turnover of the intestinal epithelium suggested to occur every 5-7 days^1^. This rapid self-renewal is facilitated by distinct proliferative epithelial populations within the crypt including active intestinal stem cells (ISC) at the crypt base, transit amplifying (TA) cells, and putative quiescent reserve ISCs (rISCs)^2, 3^. These populations have different proliferative rates, with active ISCs proliferating about once daily^1, 4^, TA cells cycling multiple times per day as they rise from the crypt and differentiate^2, 5^, and rISCs believed to be largely quiescent/slow-cycling at homeostasis yet able to increase proliferation to repopulate crypts upon injuries which ablate the more-proliferative ISCs and TA cells (e.g. irradiation, chemotherapeutics, toxins)^3, 6–16^. These distinct cycling behaviors suggest that cell cycle state may influence three core cell state decisions: survival after injury, regenerative activation after injury, or cell cycle exit toward differentiation. However, the regulatory networks that establish these different cycling states to tune regenerative ability and fate decisions are not well understood.

SRY-Box Transcription Factor 9 (SOX9) is a transcription factor known to regulate stem cell maintenance and proliferation dynamics across adult murine tissues including the intestine, colon, pancreas, and liver^17–19^. Previous work from our lab described distinct expression levels of SOX9 present across proliferative and dormant cell populations within the murine small intestinal crypt^11, 20–22^. Using a SOX9^EGFP^ reporter mouse, we demonstrated that crypt cells with high SOX9 expression (SOX9-high) are largely a quiescent or slowly-cycling population, with high SOX9 expression marking 70% of label-retaining cells within intestinal crypts^11, 23^. We further showed that subsets of SOX9-high cells co-express Ki67 after radiation during crypt regeneration, indicating that they retain or reacquire proliferative capacity under injury conditions^22^. Additional work by our group demonstrated that mice lacking intestinal SOX9 failed to regenerate following irradiation, establishing that SOX9 expression is required for reserve intestinal stem cell activity and injury-induced epithelial repair^11^. However, how SOX9 mechanistically supports rISC properties of surviving replication-dependent injury and reverting to a proliferative state is unknown.

Several studies indicate that SOX9 restrains proliferation in the crypt. SOX9 expression exhibits a graded inverse relationship with proliferative activity across intestinal crypt lineages during homeostasis, such that cells with higher endogenous SOX9 levels are associated with progressively reduced proliferation^20^ Consistent with this relationship, SOX9 loss increases crypt cell proliferation in SOX9KO mice^24, 25^ whereas forced sustained SOX9 expression suppresses proliferation in an intestinal epithelial cell line^21^. Independent evidence also links cell cycle restraint to crypt protection, as pharmacological G1 arrest increases radioresistance in the mouse intestinal crypt^26^, likely by extending the window for damage sensing, checkpoint activation, and DNA repair^27–29^. This relationship between reduced proliferation and injury survival is consistent with rISC biology, where label-retaining cells and multiple quiescent crypt populations have been shown to support regeneration after injury.^14, 23, 30–32^ Together, these findings led us to hypothesize that higher SOX9 expression supports rISC activity by reversibly elongating G1, allowing cells to survive replication-dependent injury and later re-enter a proliferative stem cell state as SOX9 levels decline.

Most studies on SOX9 function in the intestines have relied on mouse models^11, 20–22, 24, 25^, yet emerging studies show that murine and human ISCs differ at both transcriptomic and functional levels^33–35^. These differences raise questions about whether proliferative control mechanisms defined in mice are fully conserved in human ISCs and support testing ISC regulatory mechanisms in primary healthy human ISCs. To do this, we used our human monolayer model in which crypts from human donors without known GI pathology are cultured and expanded on soft collagen as proliferative monolayers consisting of stem and progenitor cells^36–38^. This monolayer system gives physiologically relevant readouts of proliferation, differentiation, and injury responses^36, 39–41^. To test whether SOX9 elongates G1 and promotes injury survival, we genetically modified primary human ISCs to tune SOX9 expression levels and quantify cell cycle dynamics, identified downstream effectors of SOX9-mediated cell cycle control, and tested whether increased SOX9 and its downstream mechanism are necessary and sufficient to confer chemoresistance to human ISCs.

## Results

### SOX9 marks proliferative human intestinal cells in vivo and in vitro

Nearly all published reports detailing expression and effects of SOX9 within the small intestine have come from mice, so we began by visualizing SOX9 expression in the human small intestinal epithelium via immunofluorescence. Consistent with murine intestines, SOX9 was expressed at various levels across cells of the proliferative crypts, and all proliferating cells expressed SOX9 **(Figure 1A)**. To lay the foundation for further studies of SOX9 using our primary ISC monolayer culture platform, we performed a similar analysis on wildtype (WT) human monolayers cultured in conditions that support ISC maintenance^38^. As monolayers expand, cells in their central region experience contact inhibition and become quiescent, establishing proliferative and non-proliferative regions **(Figure 1B)**. After three days of growth, SOX9 expression was seen throughout the proliferative region of growing monolayers, with all proliferating cells expressing SOX9 **(Figure 1B)**. SOX9 expression in all proliferative cells across human crypts and monolayers, with minimal expression seen in non-proliferative populations in both settings, indicates that SOX9 may have conserved, physiologically-relevant cell cycle regulatory effects in vivo and in vitro.

**Figure 1:**
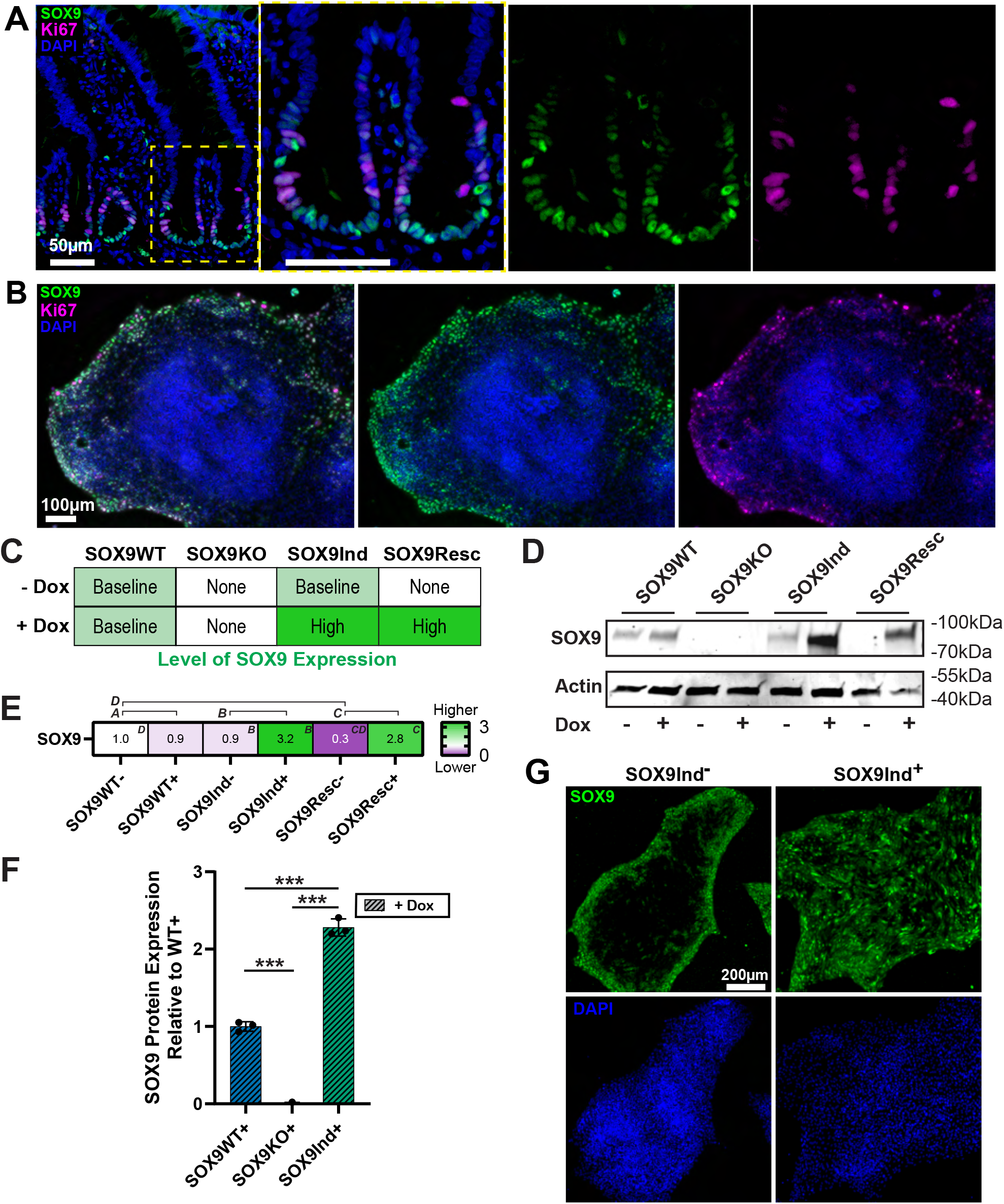
Engineering primary human ISCs to control SOX9 expression. **A)** Immunofluorescence for SOX9 (Green), Ki67 (Fuchsia), and DAPI (Blue) in human jejunal crypts. **B)** Immunofluorescence for SOX9 (Green), Ki67 (Fuchsia), and DAPI (Blue) in human jejunal monolayers cultured for three days in stem cell-promoting conditions. **C)** Schematic of SOX9 expression levels with/without Doxycycline (Dox) treatment across engineered cell lines. Dox induces SOX9 expression in SOX9Ind and SOX9Resc cells; SOX9 knockout is constitutive. **D)** Western blot demonstrating SOX9 expression differences across the cell lines depicted in (C) cultured with or without 100 ng/mL Dox for three days. **E)** Heatmap indicating transcripts normalized to SOX9WT-noDox expression levels. Green: Higher expression; Purple: lower expression. Superscript letters indicate significant differences (p<0.05 via two-tailed t-test) across pairs of conditions demarcated by the brackets on top of the heatmap. **F)** Quantification of Western blot results comparing three wells each of WT, SOX9KO, and SOX9Ind cell lines treated with 100 ng/mL Dox for 3 days. **G)** Immunofluorescence for SOX9 (Green) and DAPI (Blue) in SOX9Ind monolayers grown with (right) or without (left) 100 ng/mL Dox for three days.

### SOX9 dosage restrains proliferation

To probe SOX9 function in primary human ISCs, we genetically engineered a constitutive knockout line (SOX9KO), a line with doxycycline (Dox)-<u>ind</u>ucible SOX9 overexpression (SOX9Ind), and a <u>resc</u>ue line in which inducible SOX9 restores SOX9 in the knockout background (SOX9Resc) (**Figure 1C)**. Western blotting showed Dox did not alter SOX9 expression in WT cells, confirmed SOX9 loss in SOX9KO cells regardless of Dox treatment and in untreated SOX9Resc cells, and showed increased Sox9 expression in Dox treated SOX9Ind or SOX9Resc lines **(Figure 1D)**. Transcriptomic analysis further validated the expected SOX9 mRNA patterns across these conditions, distinguishing negligible Dox-only effects in WT cells (pair ‘A’), SOX9 induction from baseline (pair ‘B’), SOX9 rescue in the knockout background (pair ‘C’), and SOX9 loss relative to WT controls (pair ‘D’) **(Figure 1E)**. Additional Western blotting quantified SOX9 protein changes across populations, showing no expression in SOX9KO cells and SOX9 induction (SOX9Ind+Dox) increasing total levels 2.28-fold (**Figure 1F**, p<0.0001). Because our previous work shows Sox9-high cells have 3-4x more SOX9 protein expression than Sox9-low ISCs in murine crypts^20^, these induced levels appear to fall within the physiological range. Finally, SOX9 immunofluorescence confirms that SOX9 induction expands the SOX9-positive region beyond the monolayer edges seen at baseline **(Figure 1G)**.

Because SOX9 loss increases intestinal proliferation in mice^11, 24^, we tested whether SOX9 similarly regulates proliferation in human ISCs. Following a 1-hour EdU pulse, SOX9KO and untreated SOX9Resc (no Dox) cells showed visibly increased EdU incorporation compared with WT cells, whereas Dox treatment reduced EdU incorporation in SOX9Ind and SOX9Resc cells but not in WT or SOX9KO controls **(Figure 2A)**. Flow cytometry confirmed these effects; SOX9KO cells contained 45.3% more EdU-positive cells compared to WT cells (**Figure 2B**, p=0.0005), while SOX9 induction reduced EdU incorporation by 44.5% in SOX9Ind and 33.9% in SOX9Resc cells (**Figure 2C**, both p<0.0001). Dox had negligible effects in WT and SOX9KO cells (**Figure 2C**, p=0.38 and p=0.45), supporting that SOX9 drives these proliferation differences rather than off-target Dox effects. Together, these results indicate that SOX9 represses proliferation in human ISCs based on its expression level.

**Figure 2:**
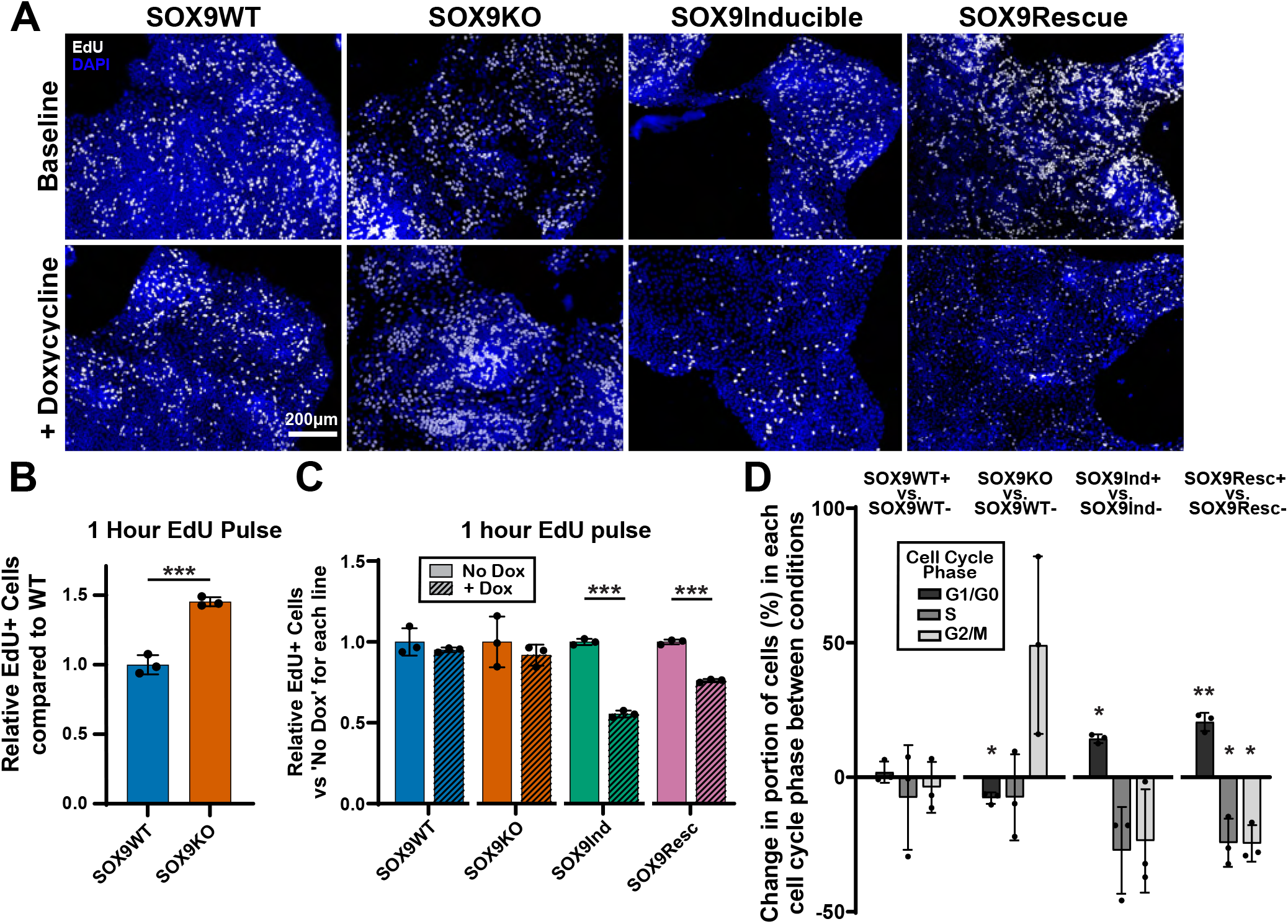
SOX9 represses proliferation in ISCs. **A)** Immunofluorescence for EdU uptake (White) in monolayers (DAPI; Blue) grown for three days +/-100 ng/mL Dox then pulsed with EdU for 1 hour prior to fixing. **B)** Flow cytometry comparing number of cells taking up EdU in WT vs SOX9KO monolayers as in (A) (*** p<0.0005 via two-tailed Student’s t-test). **C)** Flow cytometry as in (B) for WT, SOX9KO, SOX9Ind, and SOX9Resc cells grown with or without 100 ng/mL Dox for three days. For all pairs, relative uptake following Dox is shown compared to the same cell line without Dox (*** p<0.0005 via two-tailed Student’s t-test). **D)** Flow cytometry to quantify DNA content within each cell (via propidium iodide uptake) to define proportions of the total cell population within each cell cycle phase at the time of fixing. For each comparison, bars indicate the magnitude of change in proportion sizes between the top condition and the bottom condition listed. (Significance tested between values for each cell cycle phase +/-Dox, * p<0.05; ** p<0.005 via two-tailed Student’s t-test).

To determine whether SOX9 alters cell cycle phase distribution, we quantified DNA content by flow cytometry for propidium iodide to quantify proportions of cells in each cell cycle phase ^42–44^. Flow cytometry on freely growing monolayers treated with or without Dox for 3 days demonstrated that Dox alone did not alter cell cycle phase distribution in WT cells (**Figure 2D**). Compared to WT cells, SOX9KO cells showed a 49.1% increase in G2/M phase, with a small but significant 7.7% decrease in G1/G0 and no change in S phase (p=0.062, 0.0037, 0.48, respectively). Conversely, SOX9 induction shifted cells toward G0/G1 in both SOX9Ind and SOX9Resc backgrounds, increasing the G0/G1 fraction by 14.3% and 20.6%, respectively (p=0.003 and p=0.0008), while reducing cells in S phase by 27.2% and 24.4% (p=0.056 and p=0.018) and G2/M by 23.7% and 24.6% (p=0.11 and p=0.0066). These results indicate that SOX9 represses proliferation by shifting more cells to reside in the G0/G1 phases at any given time. However, DNA-content analysis cannot distinguish whether this reflects G1-elongation in actively cycling cells or increased exit into the non-proliferative G0 phase, motivating us to perform direct measurement of cell cycle phase length.

### SOX9 elongates G1 phase

To directly determine whether SOX9 regulates G1 phase length in human ISCs, we combined the SOX9 engineered lines with our PIP-H2A cell cycle reporter platform **(Figure 3A)**^36, 45^. Dually modified cells were cultured on chamber slides with pressed collagen, then live imaged to quantify cell cycle lengths in freely cycling human ISCs **(Figure 3B)**. SOX9KO;PIP-H2A cells had a 34.4% shorter G1 phase and an 11.7% shorter S-phase than WT;PIP-H2A cells, with no significant change in G2/M phase. These changes shortened total cell cycle length by 22.2%, corresponding to a 3.7 hour decrease (**Figure 3C-E**; G1/S/Total p<0.0001, G2/M phase p=0.098). Conversely, Dox-treated SOX9Ind;PIP-H2A cells had a 91.0% longer G1 phase, a 13.3% longer S phase, and a 17.4% longer G2/M phase, increasing total cell cycle length by 48.7%, or 7.3 hours (**Figure 3F-H**; G1/S/Total p<0.0001, G2/M phase p=0.0005). Dox treatment alone slightly decreased cell cycle phase lengths in WT;PIP-H2A cells (G1 p<0.0883, S p=0.0303, G2/M p=0.010, Total p=0.013), confirming that the elongation observed after SOX9 induction could not result from off-target Dox effects **(Figure 3I)**.

**Figure 3:**
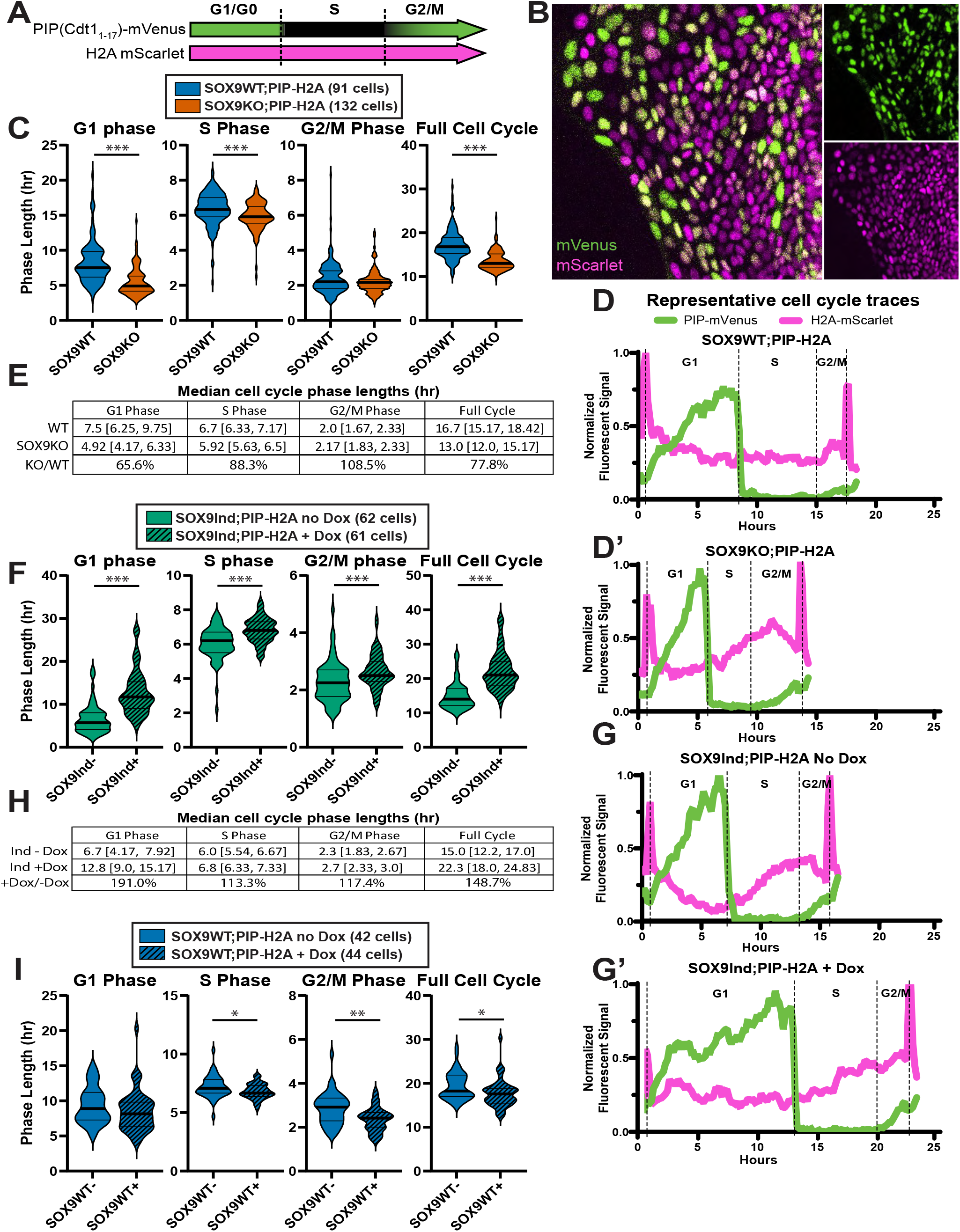
SOX9 elongates ISC cell cycle largely via lengthening G1 Phase. **A)** Schematic illustrating reporter colors that freely growing PIP-H2A cells fluoresce during each cell cycle phase. **B)** Snapshot of freely growing PIP-H2A reporter cells showing mVenus (Green) and mScarlet (Fuchsia) fluorescence in each nucleus. C) Violin plot showing lengths of each cell cycle phase and total cell cycle across WT (91 cells) and SOX9KO (132 cells). **D)** Representative live imaging tracing for both reporter colors in a PIP-H2A cell undergoing the full cycle in WT (D) or SOX9KO (D’) monolayers. **E)** Median lengths of each cell cycle phase and full cell cycle for WT and SOX9KO cells. **F)** Same as (C) comparing SOX9Ind-noDox (62 cells) and SOX9Ind+Dox (61 cells). **G)** Same as (D) comparing SOX9Ind-noDox (G) and SOX9Ind+Dox (G’). **H)** Same as (E) comparing SOX9Ind-noDox and SOX9Ind+Dox cells. **I)** Same as (C) comparing SOX9WT cells +/-Dox to determine if off-target effects of Dox alone are driving the cell cycle changes. For all violin plots, median and 1^st^+3^rd^ quartiles are marked; * p<0.05; ** p<0.005; *** p<0.0005 via two-tailed Student’s t-test or Wilcoxon-Mann-Whitney test depending on normality within the groups.

Notably, G1 accounted for most of the SOX9-dependent change in total cell cycle length. In SOX9KO ISCs, G1 phase decreased 2.58h, accounting for 69.7% of the total cell cycle shortening, even though G1 phase only accounts for 44.9% of baseline total cycle length **(Figure 3E)**. Similarly, SOX9 induction increased G1 by 6.1h, accounting for 83.6% of the 7.3h increase in total cell cycle length **(Figure 3H)**. Together, these live imaging data demonstrate that SOX9 induction regulates total cell cycle length in actively cycling human ISCs primarily by lengthening G1 phase.

### SOX9 induction does not drive differentiation

Because reduced proliferation can result from ISC differentiation, we asked whether SOX9 induction drives human ISCs to differentiate toward mature intestinal lineages. For a targeted approach, bulk RNAseq data from WT, SOX9Ind, and SOX9Resc cells grown in stem cell maintenance media with or without Dox for three days were probed for 10 canonical markers of differentiated intestinal lineages, with pairs of markers chosen for each lineage (Absorptive enterocytes = APOA4, SI; Paneth cells = PRSS2, LYS; BEST4^+^ cells = LYZ, BEST4, CA7; Enteroendocrine cells = CHGA, CHGB; Goblet cells = MUC2, SPINK4; Tuft cells = PTGS1, POU2F3)^34^ **(Figure 4A)**. Dox alone did not significantly affect any of these markers in WT cells. SOX9 induction in SOX9Ind cells decreased four markers and increased only two (SI, CHGB), and SOX9 rescue in Dox-treated SOX9Resc cells similarly decreased two markers and increased only two (CHGB, PTGS1). Importantly, these changes did not reflect coordinated activation of a differentiated lineage program. For each lineage, two independent canonical markers were assessed, and increased expression of one marker was not accompanied by increased expression of the second marker from the same lineage. Instead, the corresponding lineage marker either remained unchanged or trended downward, arguing against directed differentiation toward any mature intestinal lineage. By contrast, SOX9 loss (SOX9Resc no Dox vs WT) showed a mixed pattern, with increased expression of four markers, including both goblet cell markers, and decreased expression of two markers, both associated with tuft cells. These results suggest that SOX9 induction does not reduce proliferation by driving cells toward a defined mature intestinal lineage.

**Figure 4:**
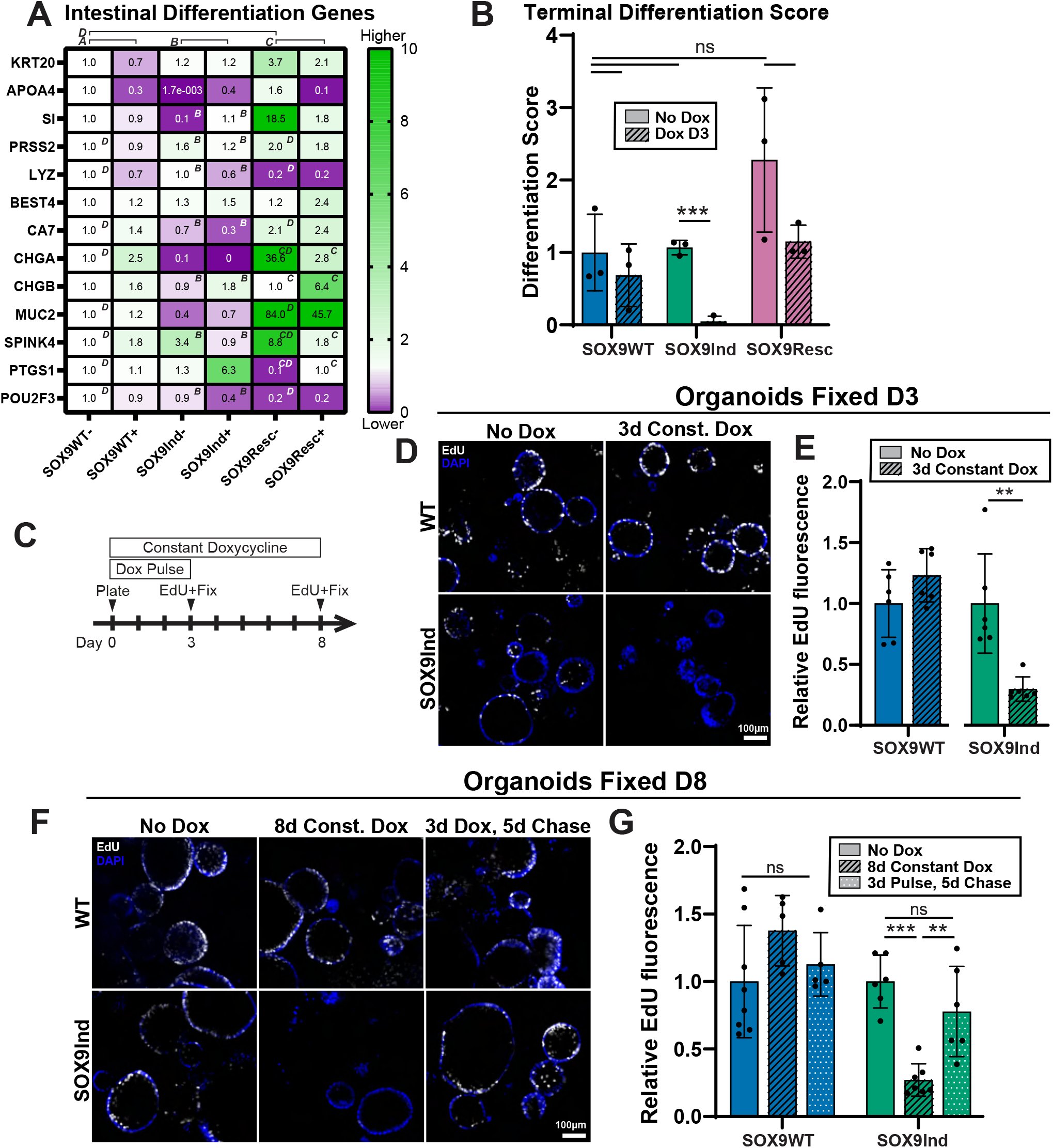
SOX9 induction reversibly represses ISC proliferation. **A)** Heatmap indicating transcript levels of canonical markers of differentiated intestinal lineages in monolayers grown in stem cell maintenance media for three days, with all values normalized to expression levels in SOX9WT-noDox cells. Green: higher expression; Purple: lower expression. Superscript letters indicate significant differences (p<0.05 via two-tailed t-test) across pairs of conditions demarcated by the brackets above the heatmap. **B)** Terminal Differentiation Score measuring how similar each monolayer condition is to monolayers grown in Differentiation Media for 0-11 days in our prior study^46^ (*** p<0.0005 via two tailed Student’s t-test). **C)** Timeline for proliferation recovery assay. **D)** Immunofluorescence for EdU uptake (White) in organoids (DAPI; Blue) grown for three days with or without 100 ng/mL Dox then pulsed with EdU for 1 hour prior to fixing. **E)** Quantification of EdU+ pixel intensity across DAPI+ pixels in images as in (D) (** p<0.005 via one-way ANOVA). **F)** Immunofluorescence for EdU uptake (White) in organoids (DAPI; Blue) grown for 8 days with or without 100 ng/mL Dox or for a 3-day Dox pulse followed by a 5-day no-Dox chase, with all conditions pulsed with EdU for 1 hour prior to fixing. **E)** Quantification of EdU+ pixel intensity across DAPI+ ixels in images as in (F) (** p<0.005, *** p<0.0005 via one-way ANOVA).

As a robust complementary approach, we used a transcriptomic differentiation score derived from our prior dataset in which ISC monolayers were followed for 0-11 days after transfer from maintenance media to differentiation media^46^. This score was based on the principal component that best captured the differentiation trajectory in our previous study. When applying this Differentiation Score to our current dataset, SOX9 deficient cells (SOX9Resc without Dox) trended toward a higher Differentiation Score despite their increased proliferation **(Figure 4B)**. This is consistent with lower/negligible SOX9 expression normally observed in TA cells and differentiated lineages.^20^ Restoring SOX9 in the SOX9Resc line returned the Differentiation Score toward WT levels, while SOX9 induction in SOX9Ind cells significantly lowered the score (p=0.0001); Dox alone had negligible effects in WT cells **(Figure 4B)**. Together, our targeted marker analysis and transcriptome-wide Differentiation Score indicate that proliferative restraint seen with increased SOX9 expression is not explained by terminal differentiation.

### Effects of SOX9 induction are reversible

SOX9-mediated control of G1 length may help explain prior irradiation phenotypes in murine intestine: SOX9 loss impairs recovery^11^ potentially by reducing survival of rISC-capable crypt cells with shortened G1 phases, and pharmacologic G1-S blockade with palbociclib enhances crypt radioresistance^26^. To test whether SOX9 induction confers rISC-associated properties in vitro, we evaluated three key functions: re-entry into a highly proliferative state, retention of stem cell activity, and survival after injuries that ablate actively cycling ISCs.

We first asked if the lower proliferation rate following SOX9 induction is reversible. WT and SOX9Ind ISCs were grown as 3-Dimensional organoids to avoid confounding effects from contact inhibition blocking proliferation in monolayers. A three-day Dox pulse reduced EdU uptake by 80.3% in SOX9Ind organoids (p=0.002), with negligible effects in Dox-treated WT controls **(Figure 4C-E)**. To determine whether this proliferative restraint could be reversed, organoids were treated with Dox for three days, Dox was washed out, and then the organoids were cultured an additional 5 days. After this chase period, EdU uptake recovered to levels similar to organoids that never received Dox (p=0.24), whereas organoids maintained in Dox for the full eight days retained an 82.9% reduction in EdU uptake **(Figure 4F-G**, p<0.0001). This demonstrates that high SOX9 places ISCs in a reversible low-proliferative state while preserving their ability to resume proliferation, a functional property associated with rISC activity.

### SOX9 induction maintains stemness

We next asked whether increasing SOX9 causes cells to cells to lose ISC identity even while they are cultured in stem cell promoting conditions. We first checked whether stem cell markers were maintained across different SOX9 levels. We analyzed 11 ISC markers defined as conserved between human and mouse ISC signatures from our intestinal atlas^34^. Dox treatment of WT ISCs did not significantly change any marker **(Figure 5A)**. Inducing SOX9 expression in SOX9Ind or SOX9Resc cells produced mixed effects across markers, with some markers decreasing, some increasing, and others remaining unchanged. In SOX9Ind cells five markers decreased, two increased, and others including the canonical ISC marker *LGR5* were unchanged. In SOX9Resc cells, seven markers decreased, while two including *LGR5* increased after SOX9 restoration **(Figure 5A)**. Thus, SOX9 induction modestly alters ISC marker expression but does not uniformly erase ISC identity and does not drive terminal differentiation. This pattern suggests that elevated SOX9 may place cells in an intermediate state: less proliferative than active ISCs, not committed to terminal differentiation, and still able to resume cycling as SOX9 levels decline.

**Figure 5:**
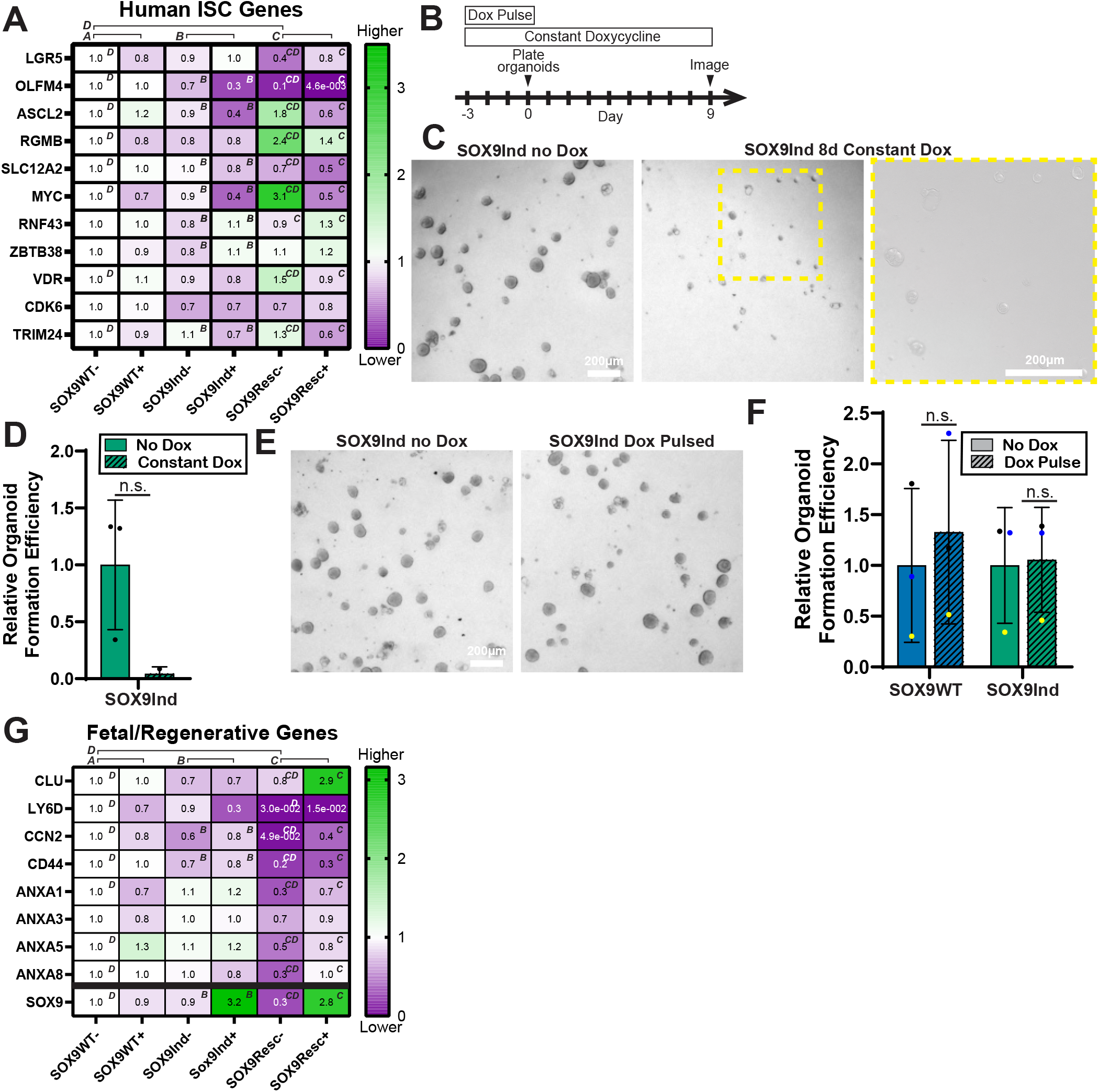
SOX9 induction maintains ISC stemness. **A)** Heatmap indicating transcript levels of Intestinal Stem Cell markers normalized to expression levels in SOX9WT-noDox cells. Green: higher expression; Purple: lower expression. Superscript letters indicate significant differences (p<0.05 via two-tailed t-test) across pairs of conditions demarcated by the brackets above the heatmap. **B)** Timeline for organoid formation efficiency assay. **C)** Brightfield imaging of organoids seeded from single SOX9Ind cells never treated with Dox or treated with 100 ng/mL Dox prior to plating and during the entire growth period. **D)** Quantifications for (C). **E)** Brightfield imaging of organoids seeded from single SOX9Ind cells never treated with Dox or treated with 100 ng/mL Dox only prior to plating but not during the organoid formation period. **F)** Quantifications for (E) normalized to no-Dox condition for each cell line. **G)** Heatmap indicating transcripts of Fetal/Regenerative markers normalized to expression levels in S X9WT-noDox cells. Green: higher expression; Purple: lower expression. Superscript letters indicate significant differences (p<0.05 via two-tailed t-test) across pairs of conditions demarcated by the brackets above the heatmap.

To determine whether SOX9 induction blocks functional stem cell activity, we tested stemness using an organoid formation efficiency assay. ISC monolayers were treated with or without Dox for three days, dissociated into single cells, and plated in Matrigel with or without Dox to quantify organoid formation **(Figure 5B)**. When SOX9Ind cells were maintained in Dox after plating, they failed to expand into organoids and instead remained as live single cells through the 9-day assay **(Figure 5C-D)**, consistent with sustained SOX9-mediated proliferative restraint. To test whether stem cell activity could recover after SOX9 levels declined, cells were treated with Dox as monolayers, dissociated to single cells, then plated into Matrigel with no additional Dox. Under these conditions where the cells could control their SOX9 expression, organoid formation by WT and SOX9Ind cells were unchanged after a 3-day Dox pulse **(Figure 5E-F)**. These results show that transient SOX9 induction does not irreversibly impair stem cell function and that SOX9-induced cells regain organoid forming capacity when SOX9 levels decline.

Because SOX9 expression is increased following multiple GI injury models^47^ and SOX9 is required for TGFB-induced fetal/regenerative reprogramming in murine intestines^48^, we next asked whether SOX9 induction itself is sufficient to drive a regenerative transcriptional state in human ISCs. SOX9 induction caused minimal changes across a panel of human fetal/regenerative genes^48^ **(Figure 5G)**. By contrast, SOX9 loss significantly decreased six of the seven genes, five of which were restored by Dox-induced SOX9 rescue. These data indicate that SOX9 is required to maintain components of a regenerative transcriptional program, but increased SOX9 alone is not sufficient to broadly induce that state. Together, these assays indicate that SOX9 induction does not drive terminal differentiation, irreversible stem cell loss, or broad regenerative reprogramming. Rather, elevated SOX9 defines a reversible low-proliferative intermediate state that preserves stem cell function as SOX9 levels decline.

### SOX9 induction protects against 5-FU chemotoxicity

A defining feature of rISC identity is survival after injuries that ablate actively cycling stem and progenitor populations^13, 23, 31^. Mouse abdominal irradiation (14-16 Gy) is commonly used to eliminate ISC and TA populations and test rISC-mediated regeneration^11, 22^. Surprisingly, even irradiation with 40 Gy was insufficient to cause any noticeable death of WT human intestinal organoids **(Figure 6A-B)**. Positive staining for H2AX confirmed that DNA damage occurred, yet organoids remained alive and proliferative three days later **(Figure 6C)**, advancing prior evidence reporting that human colonoids survive 12 Gy irradiation^49^ to show survival at even higher dosage. Thus, irradiation cannot provide a tractable injury model for testing SOX9-dependent survival in cultured primary human ISCs.

**Figure 6:**
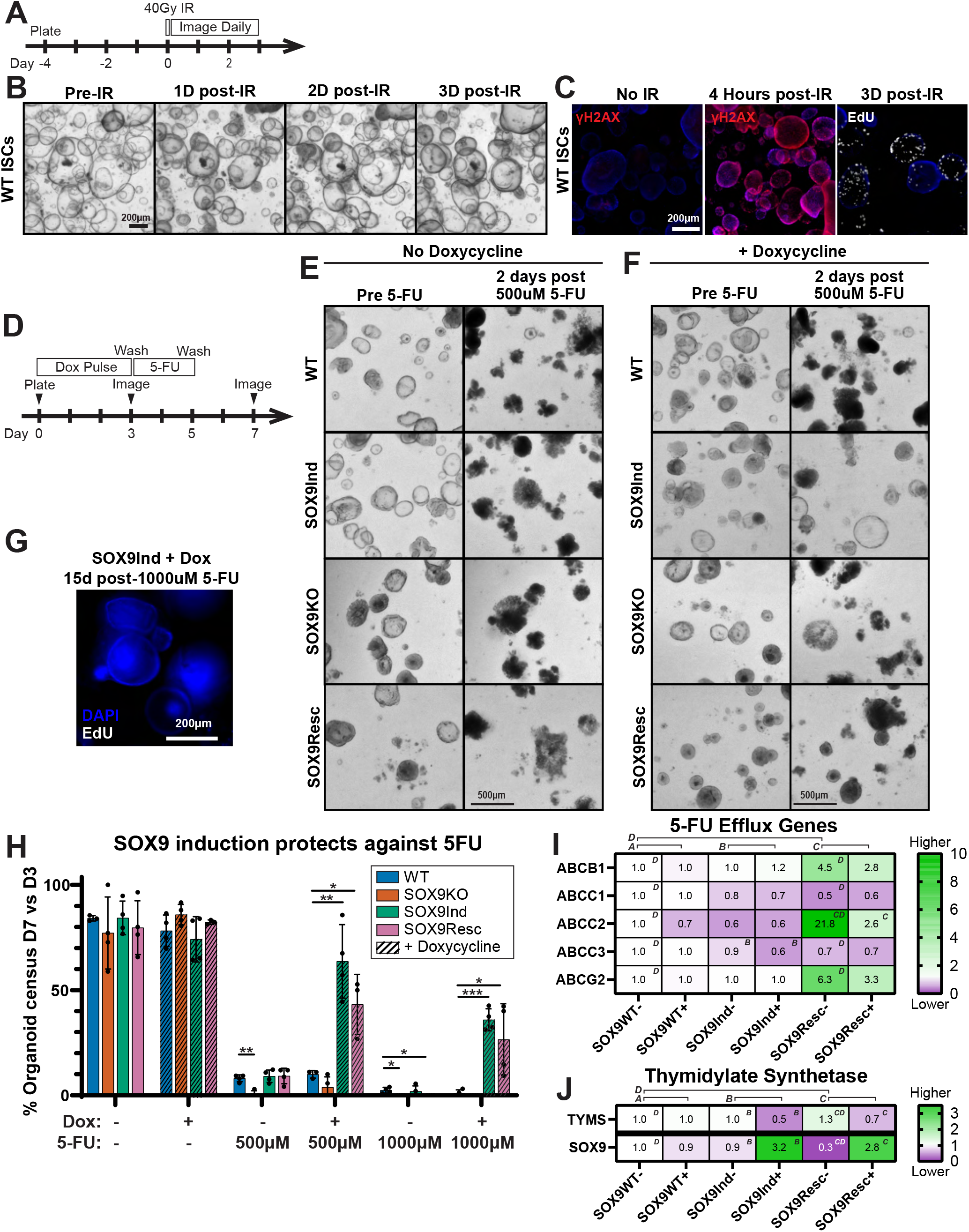
SOX9 induction grants chemoprotection against 5-FU. **A)** Timeline for organoid irradiation assay. **B)** Brightfield imaging for WT organoids daily following 40 Gy irradiation. **C)** Immunofluorescence for organoids (DAPI; blue) following irradiation stained for γH2AX (red) to denote DNA damage or EdU (white) to denote active proliferation. **D)** Timeline for 5-Fluorouracil (5-FU) assays. **E)** Brightfield imaging for organoids two days after a 48 h 5-FU treatment without Dox pre-treatment. **F)** Brightfield imaging for organoids two days after a 48 h 5-FU treatment that received three days of 100 ng/mL Dox prior to the 5-FU. **G)** Immunofluorescence for SOX9Ind+Dox organoids 15 days following 5-FU treatment (DAPI, blue; EdU, white). **H)** Quantifications for (E) and (F). (* p<0.05, ** p<0.005, *** p<0.0005 via two tailed Student’s t-test). **I-J)** Heatmaps indicating transcripts for 5-FU efflux genes (I) or thymidylate synthetase (J) normalized to SOX9WT-noDox expression levels. Green: higher expression; Purple: lower expression. Superscript letters indicate significant differences (p<0.05 via two-tailed t-test) across pairs of conditions demarcated by the brackets above the heatmap.

We therefore turned to 5-Fluorouracil (5-FU), a chemotherapeutic that causes intestinal cytotoxicity and preferentially affects proliferating cells^50^. 5-FU largely kills cycling cells by generating toxic nucleotide metabolites that are incorporated into newly synthesized DNA and RNA^51^, and prior studies suggest that slower cycling or G1 arrest can reduce 5-FU sensitivity^51, 52^. Because SOX9 induction slows ISC cycling and cell lines with different proliferation rates often harbor different 5-FU sensitivities^53–56^, 5-FU provides a replication-dependent injury model to test whether SOX9-mediated proliferative restraint promotes survival. Prior intestinal organoid studies report 5-FU cytotoxicity in the 100-1000 µM range^57, 58^, and we found that a 48h treatment with 500 µM 5-FU killed over 90% of human ISC organoids **(Figure 6D-E)**. Dox pre-treatment did not protect WT or SOX9KO organoids from 5-FU ablation (WT p=0.21; KO p=0.26) **(Figure 6F)**. By contrast, SOX9 induction increased survival 7.0-fold in SOX9Ind organoids and 4.7-fold in SOX9Resc organoids **(Figure 6F-H**, SOX9Ind p=0.0008, SOX9Resc p=0.0053). Protection was even observed after 1000 µM 5-FU treatment **(Figure 6F,H)**. Live organoids remained evident 15d post treatment, the latest time point we could assess before Matrigel dome breakdown; however, no EdU uptake was detected at this time point **(Figure 6G)** consistent with reports that 5-FU-treated cancer lines can require 2-4 weeks to restart expansion^59, 60^.

To determine whether this protection reflected canonical 5-FU resistance mechanisms, we examined genes involved in 5-FU efflux and target availability. Upregulation of efflux genes is a conserved mechanism of 5-FU resistance in colorectal cancer^52^, but these genes were either unchanged or decreased after Dox treatment in SOX9Ind or SOX9Resc cells **(Figure 6I)**. Similarly, upregulation of thymidylate synthetase, an enzymatic target of 5-FU is a well-established 5-FU resistance mechanism ^61^, yet *TYMS* decreased after SOX9 induction **(Figure 6J)**. These results suggest that SOX9 protection is not explained by increased 5-FU efflux or targets and may instead reflect changes associated with SOX9-induced elongation of the cell cycle.

### SOX9 elongates cell cycle via the INK4A pathway

SOX9 induction increased 5-FU survival, but this did not establish whether SOX9-induced cell cycle slowing itself was sufficient for protection, since SOX9 likely regulates additional ISC functions. To directly test whether reducing proliferation can protect ISCs from 5-FU, we treated organoids with Palbociclib for 48h before 5-FU treatment to promote G1 arrest. Palbociclib pre-treatment significantly reduced organoid death after both 500 µM and 1000 µM 5-FU treatments **(Figure 7A-B)**, indicating that cell cycle inhibition alone is sufficient to impart chemoprotection in human ISCs.

**Figure 7:**
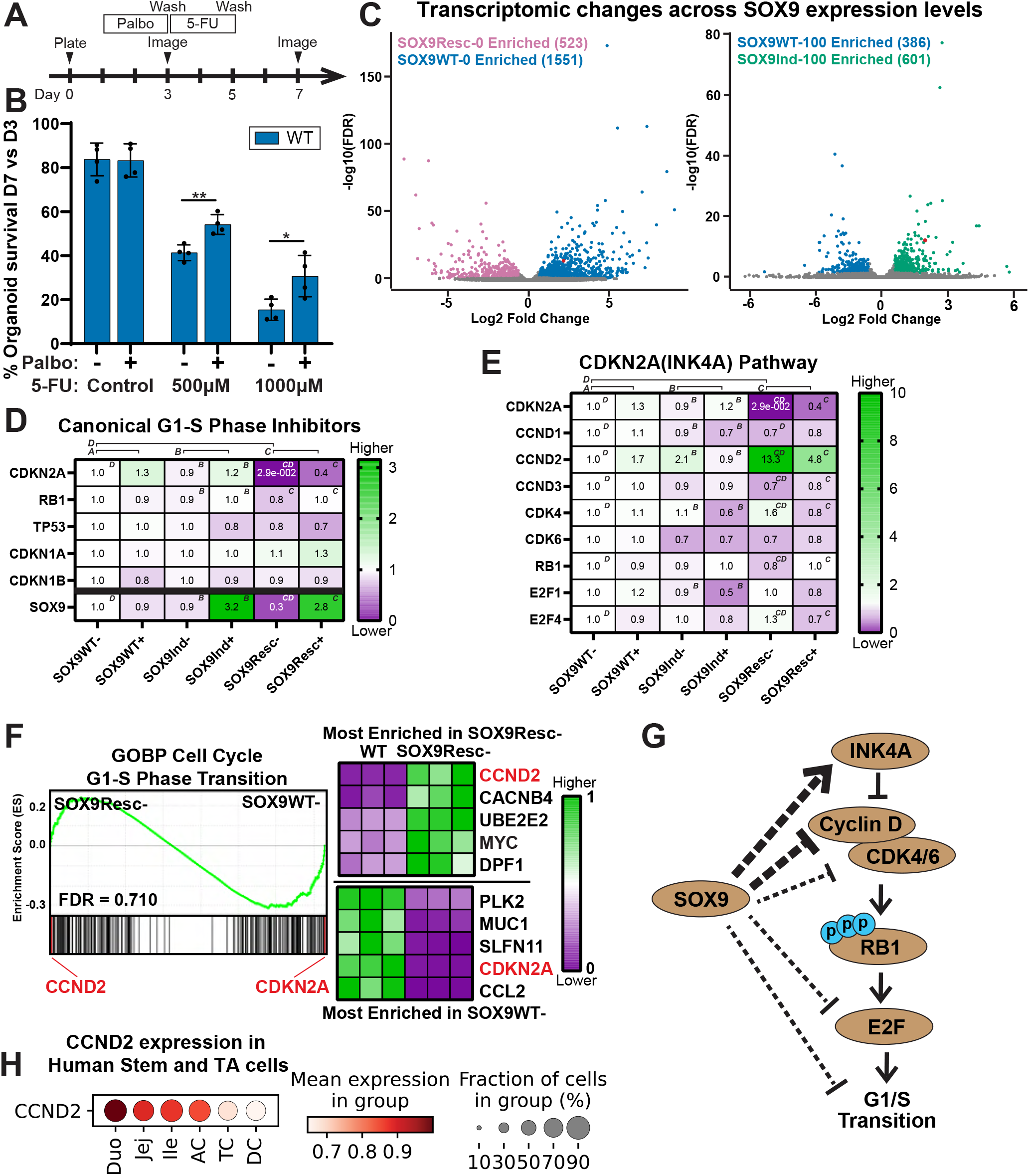
SOX9 represses cell cycle via the INK4A pathway. **A)** Timeline for Palbociclib + 5-Fluorouracil (5-FU) assay. **B)** Organoid survival quantified across WT organoids pre-treated with 48 h Palbociclib prior to 5-FU treatment (* p<0.05, ** p<0.005, via two tailed Student’s t-test). **C)** Volcano plots showing differentially expressed genes between SOX9Resc-noDox (functionally SOX9KO) and WT-noDox cells (left) or between WT+Dox and SOX9Ind+Dox (right). Gray: nonsignificant; colored: significantly different expression. False discovery rate <0.05 and a fold change of > 1.5 or < −1.5. OX9 is marked in red on both plots. **D-E)** Heatmaps indicating transcripts for canonical G1-S phase transition inhibitors (D) or major members of the INK4A-Rb Pathway (E) normalized to SOX9WT-noDox expression levels. Green: Higher expression; Purple: lower expression. Superscript letters indicate significant differences (p<0.05 via two-tailed t-test) across pairs of conditions demarcated by the brackets on top of the heatmap. **F)** Gene Set Enrichment Analysis for all members of the GOBP Cell Cycle G1-S Phase Transition gene set analyzed across SOX9Resc-(Functionally SOX9KO) and SOX9WT-cells (left). Top 5 most enriched genes shown for each cell line (right; Green: higher expression, Purple: lower expression). **G)** Schematic of the proposed mechanism of SOX9 repressing G1-S phase transition via the INK4A-Rb pathway. **H)** Dotplot showing *CCND2* expression using data from our transcriptomic atlas^34^ within ISC and TA lineages (combined) shown by region of the human intestine.

We next sought to identify the mechanism by which SOX9 elongates G1. Because SOX9 is a transcription factor and pioneer factor^62–65^ and our transcriptomic data showed broad SOX9-dependent gene-expression changes **(Figure 7C)**, we focused on expression levels of canonical regulators of the G1-S transition. Among G1-S inhibitory genes, *CDKN2A* (INK4A) showed strong SOX9-dependent positive regulation: SOX9 loss reduced CDKN2A expression by 97.1%, whereas SOX9 induction increased *CDKN2A* expression by 29.3% in SOX9Ind cells **(Figure 7D)**. Although the INK4A–Rb pathway is regulated substantially at the post-transcriptional level^66^, several downstream pathway components were also altered at the expression level with SOX9 dosage. *CCND2* (Cyclin D2), which promotes G1-S progression, increased 13-fold with SOX9 loss and decreased by 54.9% after SOX9 induction **(Figure 7D)**. *CDK4* showed a similar pattern, increasing by 57.4% with SOX9 loss and decreasing by 43.4% after SOX9 induction **(Figure 7E)**. The pro-proliferative transcription factors *E2F1* and *E2F4* were also repressed by SOX9 induction **(Figure 7E)**.

To determine whether these changes reflected broader regulation of the G1-S program, we analyzed the full GOBP Cell Cycle G1-S Phase Transition gene set comparing cells lacking SOX9 vs WT. While no enrichment for the full gene set was observed, this highlighted the substantial changes in *CDKN2A* and *CCND2* expression: *CDKN2A* was among the most decreased genes in SOX9-deficient cells while *CCND2* was the most increased gene in this pathway **(Figure 7F)**. Together, these data indicate that SOX9 regulates complementary arms of the INK4A–Rb pathway, increasing an inhibitor of G1-S progression while repressing pro-proliferative drivers, thereby providing a transcriptional mechanism for SOX9-mediated G1 elongation in human ISCs **(Figure 7G)**.

### The INK4A-Rb pathway is necessary and sufficient for 5-FU chemoprotection

To mechanistically test whether the INK4A-Rb pathway is sufficient to drive 5-FU chemoprotection, we engineered INK4A-Inducible (INK4A-Ind) ISCs in WT and SOX9KO backgrounds. INK4A induction reduced proliferation in both backgrounds, confirming that the INK4A acts downstream of SOX9 mediated cell cycle control **(Figure 8A-B)**. We titrated Dox to achieve an approximate 50% decrease in EdU uptake, matching the proliferative restraint observed after SOX9 induction **(Figure 2B)**. When these cells were challenged with 500µM 5-FU, organoids without Dox were nearly completely ablated, as observed previously **(Fig. 2B)**. By contrast, INK4A induction before 5-FU treatment increased organoid survival 3.4-fold in the WT background and 5.7-fold (p<0.0001) in the SOX9KO background **(Figure 8C-D)**, mimicking the effects of SOX9 induction. These results indicate that INK4A-mediated proliferative restraint is sufficient to protect human ISCs from 5-FU cytotoxicity.

**Figure 8:**
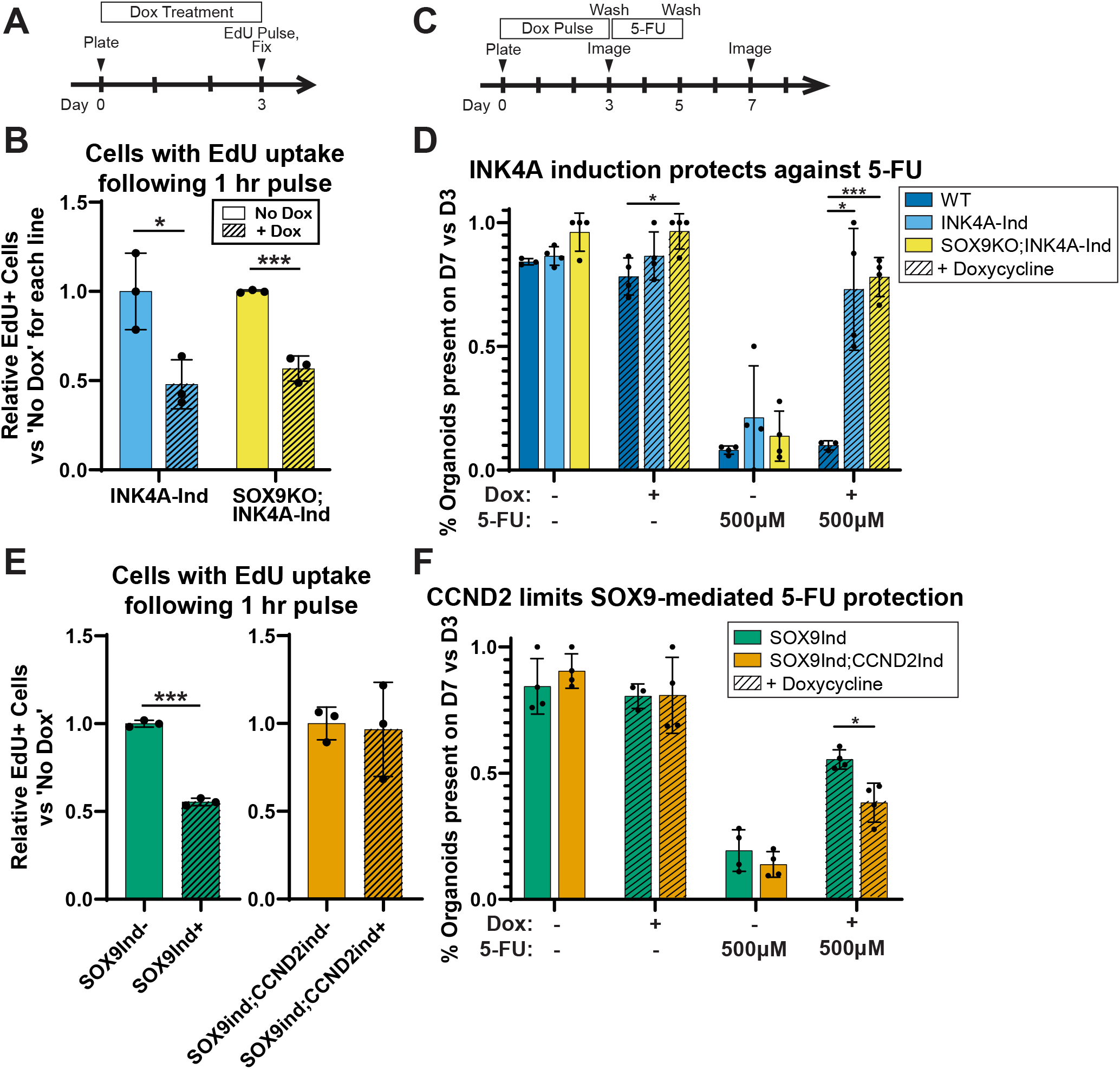
INK4A induction phenocopies the protection of SOX9 against 5-FU. **A)** Timeline for EdU uptake experiments. **B)** Flow Cytometry comparing number of cells taking up EdU in INK4A-Ind and SOX9KO;INK4A-Ind cells +/-3 days Dox treatment (25 ng/mL or 15ng/mL, respectively) (* p<0.05, *** p<0.0005 via two-tailed Student’s t-test). **C)** Timeline for 5-FU assay. **D)** Quantification of organoids surviving two days after a 48h 5-FU pulse (* p<0.05, *** p<0.0005 via two tailed Student’s t-test). **E)** Flow Cytometry comparing number of cells taking up EdU in SOX9Ind cells (left, replicated from Figure 2C) and in SOX9Ind;CCND2Ind cells +/-3 days 100ng/mL Dox treatment (*** p<0.0005 via two-tailed Student’s t-test). **F)** Quantification of organoids surviving two days after a 48h 5-FU pulse (*** p<0.0005 via two tailed Student’s t-test).

We next asked whether SOX9-mediated protection requires repression of G1-S progression. To test this, we engineered a Dox-inducible CCND2 construct in the SOX9Ind background, allowing SOX9 and Cyclin D2 to be co-induced. This approach was designed to maintain any effects SOX9 induction may have on functions besides G1-S transition while counteracting the SOX9-induced G1 phase elongation. SOX9 induction in SOX9Ind cells alone reduced EdU uptake by approximately 50% **(Figure 8E**, left panel replicating bars from Figure 2), whereas co-induction of SOX9 and CCND2 abrogated this proliferative restraint, producing similar EdU-positive fractions in Dox-treated and untreated cells **(Figure 8E).** We then tested whether this rescue of G1-S progression altered SOX9-mediated survival after 5-FU. Without Dox, SOX9Ind and SOX9Ind;CCND2Ind organoids showed comparable 5-FU-induced death, with <20% survival. SOX9 induction alone allowed 55.1% of organoids to survive, whereas co-inducing CCND2 with SOX9 repressed this protection, lowering survival to 38.1% **(Figure 8F).** Thus, CCND2 induction diminishes SOX9-mediated chemoprotection, indicating that control of the G1-S transition is a major functional component of how SOX9 protects human ISCs from 5-FU injury.

## Discussion

Dynamic cell cycle regulation is important for ISC fate decisions at homeostasis and for intestinal injury responses, yet the mechanisms that tune cell cycle length in human ISCs and connect this regulation to rISC activity remain poorly defined^67, 68^. Prior work supports a role for SOX9 in these processes: SOX9KO mice exhibit increased intestinal proliferation^24, 25^, SOX9 expression is required for epithelial recovery after irradiation-induced ISC ablation^11^, and SOX9-high crypt cells can proliferate during post-irradiation regeneration^4^. However, whether SOX9 directly regulates cell cycle dynamics, how this occurs mechanistically, and whether this regulation confers rISC-associated properties in primary human intestinal epithelium have remained unclear. Here, we show that graded SOX9 levels actively tune human ISC cell cycle length through controlling G1 phase length, creating a reversible low-proliferative state that preserves stem cell potential and promotes survival after replication-dependent injury.

A central finding of this study is that SOX9 does not simply mark a less proliferative crypt-cell state. Instead, SOX9 dosage controls human ISC cell cycle kinetics, with higher SOX9 elongating G1 and lower SOX9 accelerating cell cycle progression. This provides a mechanistic basis for prior observations that SOX9 expression is inversely associated with proliferation across crypt lineages and suggests that physiological variation in SOX9 levels may tune how crypt cells balance proliferation, restraint, and injury survival^20, 21, 24, 25^. In this model, SOX9 is not an on/off determinant of cell cycle state, but a dosage-sensitive regulator that can shift cells along a continuum of proliferative capacity.

The SOX9-high state may be best understood as a reversible competence state rather than a fixed identity. SOX9 induction restrained proliferation without broadly inducing mature lineage markers, erasing ISC identity, or activating a regenerative transcriptional program on its own. Instead, SOX9-high cells occupied an intermediate state: less proliferative than active ISCs, not committed to terminal differentiation, and capable of returning to higher proliferation and functional stem cell activity when SOX9 levels declined. This suggests that SOX9 may delay commitment while preserving responsiveness to external cues. Given that SOX9 can act as a pioneer factor^64, 65, 69, 70^ and can function through context-dependent partner proteins^71–73^, this permissive state may arise through direct transcriptional regulation, broader chromatin remodeling, or both. Defining how SOX9-high cells respond to niche, inflammatory, metabolic, and injury-associated cues may therefore provide a powerful entry point for resolving how individual crypt cells choose between renewed stem cell expansion, differentiation, and regeneration.

Mechanistically, our findings support a model in which SOX9 sets a G1-S entry threshold through the INK4A-Rb axis. Rather than acting through a single downstream target, SOX9 appears to coordinate opposing arms of the G1-S transition by increasing *CDKN2A* (INK4A) while restraining pro-proliferative outputs, including *CCND2*, *CDK4*, and E2F-family regulators. The CCND2 response is particularly notable in the context of species differences in intestinal cell cycle control. In mouse intestine, CCND1 is often emphasized as the dominant Cyclin D family member with negligible CCND2 expression seen in the proximal small intestine^74^; whereas data from our human intestinal atlas^75^ **(Figure 7H)** and from the Human Protein Atlas^76^ support CCND2 expression in proliferative human intestinal epithelial cells across the small intestine. This possibly uncovers an important difference between cell cycle maintenance in humans vs mice. Together with its strong SOX9-dependent regulation and its ability to counteract SOX9-mediated proliferative restraint during forced expression, these findings nominate CCND2 as a human ISC-relevant effector through which SOX9 levels influence cell cycle progression and injury survival. Thus, the SOX9-INK4A-Rb axis may operate as a tunable, human-relevant G1-S gate that holds SOX9-high cells in a protected, plastic state until signals favor cell cycle re-entry.

The cytoprotective effect of SOX9-mediated G1 elongation suggests a mechanism by which rISC function may emerge from normal crypt-cell heterogeneity. Rather than requiring a dedicated, permanently quiescent rISC population, rISC behavior could arise when plastic crypt cells enter a SOX9-high, G1-elongated state that reduces their exposure to S-phase-associated injury while preserving later proliferative potential. In this model, G1 acts as a protective window: cells that remain outside vulnerable rounds of DNA synthesis may be more likely to survive replication-dependent insults, avoid checkpoint-associated death, and remain available for regeneration. This framework is consistent with evidence that multiple intestinal epithelial populations can contribute to repair after injury^3, 6–16^, even though the relative SOX9 levels and regenerative contributions of these populations remain incompletely resolved. Thus, rISC activity may reflect a convergence of plasticity, cell cycle position, and injury context, with SOX9 levels helping determine which crypt cells survive the initial insult and subsequent extrinsic cues shaping whether those surviving cells re-enter proliferation, differentiate, or adopt a regenerative program.

Since irradiation induced DNA damage without ablating human intestinal organoids, 5-FU provided a tractable way to test this G1-window model in the context of replication-dependent injury. In this setting, SOX9 induction, pharmacologic G1 arrest, and INK4A induction each increased survival, whereas CCND2 induction reduced SOX9-mediated protection. Together, these experiments show that a longer G1 phase is not simply a feature of cells that survive 5-FU, but a mechanism that can directly increase survival during replication-dependent injury. Classic resistance mechanisms did not explain this phenotype, as SOX9 induction did not increase major 5-FU efflux genes and instead decreased TYMS expression. Although additional SOX9-regulated pathways may contribute to injury responses, the INK4A and CCND2 assays suggest that other pathways may be downstream of cell cycle changes. These data all support G1 elongation as a central mechanism by which SOX9-high cells gain a survival advantage during replication-dependent injury.

This work also highlights important benefits for using healthy human ISCs instead of only cancer models for studying responses to chemotherapeutics such as 5-FU. Most studies of 5-FU resistance focus on tumor cell lines, where proliferative effects are confounded by oncogenic signaling, mutations, and metabolic rewiring^53–56, 77–79^. In contrast, our engineered primary human ISC system allowed cell cycle speed to be altered within a consistent non-transformed epithelial context. The finding that G1 restraint protects healthy ISCs from 5-FU raises the possibility that normal epithelium and cancer cells can use overlapping cell cycle states for different biological ends: tissue preservation in healthy crypts versus therapeutic resistance in tumors. Our studies in healthy ISCs may even provide mechanistic insights into drug resistance in cancers, as colorectal cells with slower G1 phase can enter reversible senescence to evade 5-FU ablation^80^ and G1-lengthening factors from the tumor microenvironment are also shown to protect against 5-FU^81, 82^, showing parallels between healthy ISCs and tumor cells. CCND2 may be particularly relevant to both models, as it is implicated in GI pathologies including intestinal metaplasia and intestinal cancers^83–85^, yet in our system it functions as a SOX9-regulated G1-S effector that counteracts cytoprotective proliferative restraint in healthy ISCs. Defining how the SOX9–CCND2 axis operates in healthy versus transformed epithelium may help further distinguish physiological cytoprotective programs from pathological chemoresistance.

A limitation of our model is that it directly tests cytoprotection but not definitive post-injury regeneration. Surviving organoids could still be detected after 5-FU, but long-term analysis was limited by Matrigel dome breakdown, and EdU uptake was not observed at the latest assessable time point. Thus, while SOX9-induced cells retain stem cell potential after SOX9 decline and survive 5-FU more effectively, future lineage-tracing or live post-injury recovery studies will be required to determine whether the same surviving cells re-enter the cell cycle and regenerate epithelium after injury.

Altogether, this study identifies SOX9 dosage as a functional regulator of human ISC cell cycle length, plasticity, and injury resistance. We propose that elevated SOX9 expression establishes a reversible G1-elongated competence state through the INK4A–Rb pathway, restraining proliferation while preserving responsiveness to future cues. This model positions SOX9-mediated G1 elongation as a potential determinant of which crypt cells survive replication-dependent injury and retain reserve stem cell potential.

## Methods

### Tissue procurement and processing

Intestinal epithelial cells were harvested from human intestines following a published protocol^41^. Briefly, donor-grade human intestines were received from Honorbridge (formerly Carolina Donor Services). Jejunum was defined as the proximal half of the small intestine once the first 20cm (duodenum) was removed. Within 8 hours of organ resection, a 3×3 cm piece was excised from the middle of the jejunal length, then stored in Advanced DMEM/F12 + 10 µM Y27632 + 200 µg/mL Primocin on ice until crypt isolation. To reduce mucus, the tissue was incubated in PBS + 10 mM N-acetylcysteine for 15 min then transferred to Isolation Buffer (5.6 mM Na2HPO4, 8.0 mM KH2PO4, 96.2 mM NaCl, 1.6 mM KCl, 43.4 mM Sucrose, and 54.9mM d-sorbitol) + 2 mM EDTA + 0.5mM DTT for 30 min with gentle rocking at room temperature then vigorous shaking for 2 min. After shaking, the tissue was transferred to a new tube of Isolation Buffer + EDTA + DTT and the rocking then shaking were repeated. This was repeated six times, with supernatant for each stored on ice. Following the shakes, supernatants from each round were checked via light microscope for the presence of crypts and villi. Crypt-enriched shakes were pooled, washed in Isolation Buffer, quantified, then cultured or frozen. All monolayers and organoids for this paper were derived from jejunal crypts from one male Hispanic donor, aged 34, with no known GI pathologies. No IRB was necessary for receiving post-mortem tissue.

### Tissue culture

Tissue culture conditions are reported following Guiding Principles published in CMGH^86^. Human IECs were cultured on collagen plates prepared following a published protocol^37^. Briefly, Rat Tail Collagen I was diluted to 1 mg/mL using ice cold neutralization buffer (1x dPBS, 1 M HEPEs, 7.5% NaHCO_3_, 1 N NaOH, in DI Water), thoroughly mixed, then added to Corning Costar Cell Culture Plates (1 mL per 6w well, 500 μl per 12w well, 300 μl per 24w well). Plates were tapped until collagen evenly covered the bottom of the wells and then incubated at 37 °C for 90 min. Final collagen-coated wells were overlaid with dPBS and stored in sealed bags until use. All collagen plates throughout the study were made by an experienced specialist within the UNC Advanced Analytics Core, and monolayers were visually inspected for consistent growth phenotypes (via brightfield microscopy) across batches.

Maintenance Media was prepared as previously defined^87^. Briefly, L cells expressing transgenic WNT3A, NOGGIN, and RSPONDIN3 (ATCC CRL-3276) were cultured in Collection Media (20% Tetracycline-negative Fetal Bovine Serum, 1% Glutamax, 1%Pen/Strep, in Adv DMEM/F12) for 8 days, with media collected daily. Maintenance Media consisted of 50% conditioned Collection Media and a final concentration of 1x B-27 Supplement (without vitamin A), 10 mM Nicotinamide, 10 mM HEPES, 1X Glutamax, 1X Pen/Strep, 1.25 mM N-Acetylcysteine, 50 µg/mL Primocin, 3 µM SB202190, 50 ng/mL murine EGF, 10 nM Gastrin, 500nM A83-01, and 10 nM Prostaglandin E2. Conditioned media and Maintenance Media were prepared on-site by the UNC Advanced Analytics Core and used within 2 weeks of preparation. Fresh crypts were plated with 200 mg/mL Primocin, 200 mg/mL Gentamycin, and 0.5 mg/mL amphotericin B for the first week to avoid growth of contaminating species. Cells were grown in a humidified incubator at 37 °C with 5% CO_2._

Monolayers were passaged every 4-7 days, generally at ratios of 1:3 or 1:4. For passaging cells from a 12-well or 6-well collagen plate, the collagen patty and 1 mL of media from each well were incubated in a 37 °C water bath with 150 ul of 5,000 U/mL collagenase IV until collagen completely dissolved, centrifuged at 500 xG for 2 min, washed in PBS, then digested in 1 mL TrypLE Express + 10uM Y27632 for 5 min in a 37 °C water bath. Cells were dissociated by triturating with a P1000 pipet tip then quenched with an equal volume of 10% FBS. Cells were then resuspended in Maintenance Media + 10uM Y27632 and replated. Media was changed at least every other day.

Cells were cryopreserved in liquid nitrogen. To freeze cells, monolayers went through the same collagenase and TrypLE procedure as for passaging, except the final cell pellet was resuspended in ice-cold freeze media (60% SI Maintenance Media + 30% Tet-negative FBS + 10% dimethyl sulfoxide) and transferred to labeled cryovials. Cells were immediately placed into a −80 °C freezer within a closed Styrofoam cooler. The next day cells were transferred to a vapor phase liquid nitrogen cooler. For thawing, cryovials were held in a 37 °C water bath until ∼1/2 of the ice had melted, then diluted into 8 mL of Advanced DMEM + 10% FBS. Cells were then centrifuged and plated as normal. The act of cryopreserving and thawing was counted as a passage when tracking total passage number.

Organoids were grown in droplets of Growth Factor Reduced Matrigel with phenol red. Upon receiving, all Matrigel was diluted to 8mg/ml using Advanced DMEM/F12 prior to freezing aliquots. Matrigel stock and aliquots were thawed completely on ice before use, and aliquots never went through more than one additional freeze/thaw cycle before using or discarding. All cells were regularly maintained as monolayers, only plated into Matrigel 1-3 passages before experimentation when organoids were needed. When plating from monolayers, the passaging procedure above was followed, with final cells resuspended in Matrigel and plated instead of plating into collagen plates. When passaging organoids, Matrigel droplets and media were collected into a tube, slowly drawn through a 28G syringe three times to break up organoids and dissociate Matrigel, then centrifuged at 500 G, aspirated, resuspended in Matrigel, and 5-15ul patties plated in 48w or 96w plates. Plates were inverted and incubated at 37 °C for 20 min for the Matrigel to solidify, then wells were overlaid with 200 or 100 μL of MM, respectively.

Treatments were administered at various timepoints, with timelines provided for each experiment. For induction of target genes, SOX9Ind and CCND2Ind cells were treated with 100ng/mL, INK4A-Ind at 25 ng/mL, and SOX9KO;INK4A-Ind at 15ng/mL Doxycycline Hyclate. To prepare Dox, powder stock was first dissolved in ddH2O and then diluted into maintenance media for treatment. Palbociclib stock powder was first dissolved into DMSO and then administered at a final concentration of 5 μM for 48 h. For 5-FU assays, organoids were thinly plated into 5 μL Matrigel patties in four 48w wells per condition.5-FU stock powder was first dissolved in DMSO to a stock concentration of 400mM and then administered at final concentrations of 500-1000 μM for 48h. 5-FU aliquots were stored at −80 °C for up to 3 months before use. For each experiment, all replicates and internal controls were completed using a single batch/lot of media, Matrigel, or collagen plates whenever possible.

Most cellular experiments were conducted in cells passaged less than 20 times. However, due to the sequential mutagenesis and selection processes needed to attain some of the lines with multiple genetic perturbations, cells up to passage 40 were used in a few experiments. Passage-matched genetic controls were used when possible.

### Genetic engineering

All genetic engineering was completed following a previously described protocol^41^. Briefly, DNA segments of interest were isolated using restriction enzymes or amplified using CloneAmp HiFi PCR Premix. Plasmids were generated using an In-Fusion HD Cloning Kit then isolated from bacterial stocks using a QIAGEN HiSpeed Maxi kit. Plasmids were obtained from Addgene: pPIGA-PHD was a gift from Linzhao Cheng (Addgene plasmid #26778; http://n2t.net/addgene:26778; RRID:Addgene_26778)^88^. sg resistant gamma-tubulin was a gift from Maria Alvarado-Kristensson (Addgene plasmid #104433; http://n2t.net/addgene:104433; RRID:Addgene_104433)^89^. pCMV p16 INK4A was a gift from Bob Weinberg (Addgene plasmid #10916; http://n2t.net/addgene:10916; RRID:Addgene_10916)^90^. AAVS1-Puro XLone-CCND2 was a gift from Xiaoping Bao (Addgene plasmid #179845; http://n2t.net/addgene:179845; RRID:Addgene_179845)^91^. For CRISPR/Cas9 transfections, crRNA and tracrRNA oligos were purchased from IDT and resuspended at 100 pmol/mL. 2 µL (200 pmol) of each of gRNA and tracRNA were combined with 2 µL of 5X annealing buffer and 4 µL nuclease-free dH2O and annealed in a thermocycler (95 °C – 5 min, 95°C to78°C at −2 °C/s, 78 °C – 10 min, 78 °C to 25 °C at −0.1 °C /s, 25 °C – 5 min), then transferred to ice. 3 µL (60 pmol) of this mix was added to 2 µL TrueCut Cas9v2 (approximately 60 pmol), incubated at room temperature for 15 min, then stored on ice until use.

All constructs in this study were made from the same stock of WT human ISCs. Plasmids were electroporated into human cells using the Neon Transfection System 100 µL Kit. Dissociated cells were resuspended in 100 µL Neon Buffer R at 10,000 cells/µL with 6 µg of the plasmid of interest. For PIP-H2A and all inducible constructs, Super PiggyBac Transposase Expression Vector was added at 5 ng/µl. For SOX9KO, cells were resuspended with 60 pmol Cas9 + 60 pmol annealed crRNA:trac:RNA. Cells were electroporated using Neon preset #5 (1,700 V, 1 pulse, 20 ms) then immediately added to a six-well collagen plate with 3 mL Maintenance Media + 1:1000 Y27632. Around 4-7 days post transfection, colonies were selected using their respective antibiotics (Blasticidin 10 µg/mL, G418 200 µg/mL, hygromycin 200 µg/mL) for 4-8 days. For SOX9KO cells, clones were then isolated by digesting the collagen patty with 5,000 U/mL collagenase IV at 100 µL/mL at 37 °C for 25 min. Colonies were washed with dPBS, then individual colonies were picked using a 20 µl pipet over a light microscope and placed into individual wells of a 48-well collagen plate with 300 µL Maintenance Media + 1:1000 Y27632. SOX9KO lines were validated using qPCR, ultimately selecting a clone with >99% loss of SOX9 mRNA. All Inducible lines are non-clonal lines of cells following antibiotic selection, validated using immunofluorescence for the protein of interest. When possible, fluorescence activated cell sorting for the driven reporter gene (eGFP for SOX9Ind or mCherry for CCND2ind) was used to further purify positive cells. Across figures, colorblind accessible palettes were chosen to consistently represent each cell line^92^.

### Immunofluorescence

Fresh human intestinal tissue was fixed in 4% paraformaldehyde (PFA) overnight at 4 °C then transferred to 70% ethanol the next day. Tissues underwent paraffin processing and then sectioning and placement onto glass slides for downstream analysis. For staining, sections underwent routine deparaffinization and rehydration using Histoclear then an ethanol gradient. Slides were then washed with running water, permeabilized with 0.3% Triton X-100, blocked using 3% BSA in PBS, then primary antibodies were added in 3% BSA overnight at 4 °C. The following day, slides were washed 3x with PBS, secondary antibody and DAPI were added in 3% BSA for 1 hr at room temperature, then slides were washed in plenty of PBS and mounted using ProLong Gold antifade reagent. Sections were imaged on a BZ-X800 fluorescent microscope using a 20x objective.

Stem cell monolayers were fixed within their wells for 20 min with 4% PFA at room temperature then washed in dPBS and stored at 4 °C until staining. For staining, cells were permeabilized with 0.5% Triton X-100, washed with 0.75% glycine in PBS to quench free formaldehyde, blocked using 3% BSA in PBS, then primary antibodies were added in 3% BSA overnight at 4 °C. The following day, cells were washed 3x with Wash Solution (0.1% BSA, 0.2% Triton X-100, 0.05% Tween-20 in PBS), secondary antibody and DAPI were added in 3% BSA for 1 hr at room temperature, then cells were washed in plenty of PBS and stored at 4 °C in the dark until imaging. Plated cells were imaged on a Keyence BZ-X800 fluorescent microscope then analyzed with BZ-X800 analyzer software or FIJI software^93^. To visualize 5-ethynyl-2’-deoxyuridine (EdU) uptake, growing cells were pulsed with 10 µM EdU for 1 hr then fixed as above. Following permeabilization, cells were stained using EdU Reaction Buffer (4 mM CuSO_4_, 2 µM Sulfo-CY5-azide, 0.2 M Ascorbic Acid, in PBS) for one hour at room temperature protected from light, then washed in PBS and stained for other antibodies, if necessary.

For quantifying EdU in organoids to test if the effects of SOX9 on proliferation are reversible, full organoids were fixed within their Matrigel patties using warm 4% PFA for 20 minutes. EdU and DAPI were stained on these organoids using the same protocol as for monolayers. Organoids were then imaged on a Zeiss LSM900 confocal microscope using a 20x objective. Two non-overlapping images were taken per well, with three wells tested per condition. The images were then analyzed using Fiji software to determine the mean EdU fluorescence across all DAPI^+^ pixels: nuclei that were in focus for each image were segmented using the Otsu thresholding filter on Fiji, and then these regions were analyzed for mean EdU content. Mean values were graphed as a bar chart using GraphPad Prism, with significance tested via Student’s t-test for each condition.

### Western Blotting

Prior to harvesting protein for Western blotting, collagen plated cells were stringently washed to avoid carryover of collagen protein to the final protein collection. Monolayers and 1mL of their culture media were moved to a tube and incubated with 220 ul of 5,000 U/mL collagenase IV (50% more than for normal passaging) for a full 5 minutes past when the last trace of collagen visibly disappeared, generally ∼30min. Cells were then washed 3x in dPBS to remove collagen prior to digestion in RIPA Buffer (150mM NaCl, 50mM Tris-HCl, 1% TritonX-1, 0.5% Sodium Deoxycholate, 0.1% SDS in ddH_2_O) with Phosphatase and Protease Inhibitors added. Cells were vortexed, left in RIPA on ice for 30 min, and vortexed once more before being stored at −80 °C. Prior to measuring protein concentrations, lysates were centrifuged at top speed in a bench-top microcentrifuge in a 4 °C cold room, then supernatants transferred to a second ice-cold tube to remove non-soluble cell matter. Protein concentrations were quantified using the Pierce BCA Protein Analysis Kit following the manufacturer’s protocol, with imaging done on a CLARIOstar Plus Plate Reader.

To prepare for gel electrophoresis, 20 μg protein in RIPA buffer were made into a solution with 10% 2-mercaptoethanol and 25% 4x Laemmli Blue then heated to 95 °C for 5 minutes. Proteins were added to a Bio-Rad Mini-PROTEAN 4-15% TGX Gel in a Mini-PROTEAN Tetra Vertical Electrophoresis Cell for Precast Gels in Running Buffer (25 mM Tris, 190 mM glycine, 0.1% SDS) alongside a lane loaded with PageRuler Plus Pre-Stained Ladder. Electrophoresis was performed, starting at 50 V for 30 min and then 150 V until blue bands reached the bottom of the gel. Proteins were transferred from the gel to a methanol-activated PVDF membrane in transfer buffer (25 mM Tris, 190 mM glycine, 20% methanol) at 30V overnight.

The following day, the membrane was removed, washed in TBS (20 mM Tris pH 7.5, 150 mM NaCl) with 0.1% Tween20 (TBST), blocked for one hour, then incubated overnight with primary antibodies overnight at 4 °C in LICOR Intercept Blocking Buffer. The next day antibodies were washed off using TBST and then blocked again and secondary antibodies added for one hour at room temperature. The gel was washed with TBS and then imaged on a LI-COR Odyssey Infrared Imaging System. Bands were analyzed via Image Studio Software (Version 5.2) using the LI-COR One-Color Protein Marker (928-40000). To quantify SOX9 expression levels, N=3 wells grown side-by-side were harvested, and intensities were normalized to Actin bands to account for differential sample loading.

### RNA Extraction and analysis

For RNA extraction, monolayers were treated with collagenase IV at 150 µL/mL at 37 °C for ∼25 min. Cells were then washed with dPBS, pelleted, then resuspended in 300 µL Lysis buffer from an RNAqueous Micro Total RNA Isolation Kit. RNA was isolated using the kit following the manufacturer’s instructions. Concentration of the eluted RNA was measured on a Qubit 3 Fluorometer using a Qubit™ RNA High Sensitivity (HS) Assay Kit.

For Bulk RNA sequencing, RNA was prepared as above from N=3 wells each for WT, SOX9Ind, or SOX9Resc cells plated as monolayers for three days with or without 100ng/mL Dox treatment. RNA quality was validated using the Agilent 2100 Bioanalyzer, with all having an RNA Integrity Number > 8. Integrated fluidic circuits for gene expression and genotyping analysis were prepared using the Advanta RNA-Seq NGS Library Prep Kit for the Fluidigm Juno. Sequencing was done via NovaSeq paired end run, with 1 mismatch used in demultiplexing indexes with bcl2fastq version 2.20.0. Gene level expression was obtained through pseudo alignment of reads to human genome GRCh38 using Kallisto^94^. PC and sample correlation analysis were done with Bioconductor packages Biobase, cluster, and qvalue^95^. Expression values for plotting were obtained by TMM normalization across all samples using EdgeR package, and differential expression across samples was calculated from raw counts with DESeq2^96, 97^.

For designing a Terminal Differentiation Score, previously published bulk RNA sequencing data from monolayers treated with Differentiation Media (DM) was used^98^. A Principal Component Analysis showed that PC2 perfectly lined up all differentiating samples based on number of days in DM, with day 2 DM on bottom and day 11 DM on top. Genes from PC2 that had at least 500 reads (to avoid noise) were assembled into a Differentiation Score, with the value for untreated WT cells normalized to 1.

### Live imaging

Live imaging of PIP-H2A cells was completed exactly as in our introductory publications for the cells^36^ and the platform used to grow and image them^45^. For clarity, the methods from these papers are repeated here with minor updates:

To reproducibly coat an Ibidi µ-Slide 4 Well chamber slide with 0.3 mm collagen, a custom collagen press^45^ was created and fabricated using chlorinated polyethylene and an Ultimaker S3 3D printer. The feet of the press were affixed with coverslip glass using Glass Glue, and Rain-X was applied following the manufacturer’s guidelines to allow for removing the press without pulling up the thin collagen layer. Collagen was prepared as above. Each chamber received 250 μL of liquid collagen then tapped to have the collagen completely cover the base of the wells, the press was immediately positioned on top, then the chamber slide with the press was moved to a 37 °C incubator for 90 min for the collagen to solidify. Afterward, the presses were gently detached and dPBS was added to cover the collagen. Prepared slides were stored at room temperature in zip-top bags until use.

PIP-H2A fluorescent reporter cells were seeded onto collagen-coated chamber slides 24-48 hours before imaging, and growth media was changed the morning of the imaging session. Live imaging was performed on an Andor Dragonfly Spinning Disk Confocal Microscope mounted on a Leica DMi8 microscope stand, using a Leica HC PL APO 20x/0.75 LWD air objective with pinhole size set to 40 μm. The camera was an Andor iXon Life 888 EM-CCD, with Electron Multiplying Gain set to 150, Horizontal Shift Speed of 10 MHz – 16 bit, 2X Pre Amp Gain, 2.2 μs Vertical Shift Speed, Normal Vertical Clock Voltage and binned 2×2. Live imaging was started on fairly small colonies to allow room for growth and minimizing contact inhibition during the imaging timeframe.

To image the mScarlet gene in the PIP-H2A construct, a 561 nm laser was used for excitation, and light was collected with a 593/43 Semrock emission filter using a HC Fluotar L 25X/0.95 W 0.17 VISIR objective. For mVenus imaging, a 514 nm laser was used for excitation, and light was collected with a 538/20 Semrock emission filter. Images had 512 × 512 pixels and 1.00 μm pixel size. At each position, Z stacks were acquired using a piezo Z stage with 5 μm intervals spanning 75 μm (16 steps). 2×2 montages were acquired at each position. Each Z stack in the 2×2 montage overlapped its neighboring Z stacks by 10%. Two locations in each well were acquired per experiment. 2×2 montage Z stacks at each location were acquired every 10 minutes for 48 hours. All images directly compared to each other were acquired using the same settings. Typical settings were 600 ms exposures with 5% laser power for mScarlet, and 200 ms exposures with 1% laser power for mVenus. Temperature was maintained at 37 °C with an Okolab microscope enclosure, with continuous monitoring and feedback. 5% CO2 was warmed to 37 °C in the enclosure and humidified before being delivered to an enclosed stage top holder that contained the sample. To minimize evaporation during the experiment, 8 caps from 15mL falcon tubes were filled with water and placed surrounding the sample inside the stage-top sample holder.

### Live imaging analysis

Live image analysis for PIP-H2A cells was completed exactly as in our introductory publication^36^. For clarity, the methods from these papers are repeated here with minor updates:

Images were max projected for maximum intensity then stitched using FIJI software, with 10% overlap between quadrants. To maximize nuclear signal for tracking, the mScarlet and mVenus channel intensities were added together using FIJI to form a new channel. This summed and stitched channel was imported to CellPose2 for segmentation using our automated Macro^36^. To train the CellPose2 model, seven individual frames were used including individual and stitched positions. The “nuclei” option from the model zoo was chosen, with average nuclei diameter set to ‘12’. For training, the model predicted nuclei segmentation then a researcher would correct the results. Initial and final nuclei counts were recorded for each training session. Upon reaching >95% accuracy for three consecutive rounds, the model was considered trained. A Jupyter Notebook script was used to run the CellPose2 analysis on all 488 frames of the time-lapse image individually and save the resulting masks as a single TIFF file with 488 frames^36^. Nuclei were tracked across frames using the Fiji Trackmate Plugin. To begin, a four-channel TIFF was made combining C1: mScarlet, C2: mVenus, C3: C1+C2, C4: segmentation from Trackmate across timepoints, then imported into Trackmate. No cropping was chosen, and the Label Image Detector function was chosen using the segmentation masks in channel 4 to define nuclei. No initial thresholding or filtering was done prior to defining tracks. The “LAP Tracker” was chosen, with specifications set to 10um max distance, Gap close = 7.0, and 0 gaps allowed. After tracks were defined, tracks with total duration <480 min were filtered out to avoid fragments, as that was shorter than the shortest cell cycle recorded using the PIP-FUCCI reporter^36^. To choose tracks of nuclei which underwent a full cell cycle within the viewing period, Track name, frame, and Max Intensity of C1 values were all exported to Microsoft Excel then filtered for tracks which had at least one point where slope > |300|, following observations of dynamically brighter mScarlet signal during cytokinesis **(Figure 3D)**. Candidate tracks were monitored by eye to ascertain that the track followed a single nucleus through two splitting events, then raw data for the track was copied to an Analysis Workbook in Microsoft Excel which presents graphs of normalized mVenus and mScarlet fluorescence^36^. The G1-S transition was defined as the point after mVenus signal dropped over 50%. The S-G2 transition was defined as the point when the mVenus signal began rising above the lowest maintained level again. Cytokinesis was defined by acute peaks in mScarlet intensity. Data for a full imaging run for PIP-H2A including stitched and max-projected time lapses for mVenus and mCherry data, channel 3 resulting from adding both signals together, and the results of CellPose2 segmentation over the full timelapse, as well as the file resulting from TrackMate analysis of the timelapse imaging to so nucleus tracking, are available on Zenodo at DOI: 10.5281/zenodo.11506667

### Statistical analysis

A power analysis was run to define adequate N for PIP-H2A live imaging experiments was performed following preliminary data received comparing WT vs SOX9KO ISCs, where the two groups exhibited mean G1 cell cycle phase lengths of 8.436 and 5.796 hours, respectively with a standard deviation of approximately 3.7 within each group. To achieve robust and reliable results, the power analysis aimed for a significance level (α) of 0.05 and a power of 0.95, which is stringent enough to detect even modest effects. A priori analysis was conducted using G*Power version 3.1.9.7 to calculate a minimum sample size of 55 per group. This sample size ensures a high probability of correctly rejecting the null hypothesis, should a true effect exist, thereby minimizing the risk of Type II errors. Analyzed tracks were compiled from 3 separate colonies images across two separate wells of a chamber slide to account for locational/well-specific variability.

To compare cell cycle phase lengths, normality was first tested using the Shapiro Wilk test for each cell cycle phase of each population. Comparisons in which both populations showed normality were analyzed using a student’s t-test. Comparisons with at least one group not showing normality were analyzed using a two-tailed Wilcoxon-Mann-Whitney. * indicates p<0.05, ** indicates p<0.005, *** indicates p<0.0005. Final graphs and significance analyses were made using GraphPad Prism 9. Data is shown as violin plots for each cell cycle phase and for total cell cycle, with median values shown. Median values and quartile ranges provided were calculated using Excel.

For Volcano Plots, differential expression analyses were conducted using the edgeR package (V4.4.2) in R (V4.4.1). Genes with a false discovery rate <0.05 and a fold change of > 1.5 or < −1.5 were considered statistically significant. Volcano plots were generated using the ggplot2 package (V3.5.1).

When comparing experiments with biological N=3, a Shapiro-Wilks test was run to assess normality in a small sample size. If normal, a Student’s T-test was run, otherwise a Mann-Whitney (Wilcoxon) test was performed.

On heatmaps, pairs with p<0.05 via Two-tailed Student’s t-tests are shown with letter superscripts, with the following comparisons tested: A (SOX9WT-vs SOX9WT+), B (SOX9Ind-vs SOX9Ind+), C (SOX9Resc-vs SOX9Resc+), D (SOX9WT-vs SOX9Resc-). To best visualize differences across genes with moderate changes, the color gradient spectrum was set to maximum saturation at 10-fold increased expression.

### Single cell dissociation and Flow Cytometry

To dissociate monolayers to single cells, collagen patties were digested and cells washed as described above, with an added incubation in dPBS for 10 min at 37 °C following the collagenase step. Following the 5 min digestion in TrypLE at 37 °C, cells were triturated with a P1000 pipet 10x, drawn up and slowly expelled from a 28G syringe 6x to dissociate to single cells, then quenched with 10% FBS in Advanced DMEM/F12. Single cells were passed through a 30µm filter, and pelleted.

To quantify 5-ethynyl-2’-deoxyuridine (EdU) uptake using flow cytometry, cells were thinly plated in three wells of a 6-well plate per condition and cultured +/-Dox side-by-side. Growing cells were pulsed with 10 µM EdU for 1 hr prior to dissociation, then dissociated single cells were fixed in 6 mL ice cold 4% PFA (added while vortexing) for 20 min at 4 °C with heavy rocking. Fixed cells were washed and stored in Advanced DMEM/F12 at 4 °C. Following permeabilization, cells were stained using EdU Reaction Buffer (4 mM CuSO4, 2 µM Sulfo-CY5-azide, 0.2 M Ascorbic Acid, in PBS) for one hour at room temperature protected from light. Cells were then washed in dPBS then counted. Flow cytometry was run on a Sony SH800ZF cell sorter, with a quadrant gated directly above the no-fluorescence negatives demarcating the threshold above which all cells are considered EdU-positive.

For DNA content analysis, cells were thinly plated in three wells of a 6-well plate per condition and cultured +/-Dox side-by-side. After three days, cells were dissociated as above then fixed in 10 mL ice cold 70% ethanol for 30min at 4 °C with heavy rocking. Fixed cells were washed and stored in Advanced DMEM/F12 at 4 °C. Cells were treated with 10 µg/mL ribonuclease at 37 °C for 20 min, washed in Advanced DMEM/F12, then 6 µg/mL propidium iodide was added 10 min prior to flow cytometry. Flow cytometry was run on a Sony SH800ZF cell sorter. For cell cycle analysis via propidium iodide, damaged cells were gated against using off-target fluorescence from the unused APC channel, then propidium iodide levels in single cells were used to calculate cell cycle via the Dean-Jett-Fox Model on FlowJo v10.7.1.

### Organoid Formation and Survival Assays

For Organoid Formation Efficiency assays, cells were first plated as monolayers and treated with or without 100 ng/mL Dox for three days. Monolayers were then dissociated to single cells following the same protocol as for flow cytometry above. To best replicate common conditions in literature, cells were grown in Intesticult Organoid Growth Medium (Human) for a full passage prior to dissociation and throughout the organoid formation culture period. Live single cells were quantified on a hemacytometer with Trypan Blue, then cells were pelleted and resuspended in Matrigel. 600 live single cells were plated in one 6 µL Matrigel droplet per well on a 48w culture plate. Droplets were inverted and allowed to solidify at 37 °C for 20 minutes, then 250 µL Intesticult + Y27632 was added with or without 100 ng/mL Dox. Organoids were grown for eight days with Intesticult changed every three days; Y27632 was removed after the first media change. On the eighth day, media was replaced with warm 4% paraformaldehyde for 25 min to fix the organoids, then plates were scanned on a Keyence BZ-X800 Imager using bright field microscopy on a 4x objective. Live organoids were defined as having a smooth and tight outer barrier and being at least 4x larger than a live single cell. Organoids were counted by hand using the Cell Counter feature on Fiji software. Seven wells were counted for each condition across three separate experiments (21 total wells per condition). Data is shown as the average count for each condition across experiments relative to levels in WT-no Dox and SOX9Ind-no Dox controls, respectively. Statistical differences were calculated using two tailed Student’s T-tests across all 21 replicate wells for each condition.

For 5-FU organoid survival assays, organoids were grown in Maintenance media throughout the full assays. Y27632 was added during the first media overlay upon initial plating but then not included in the first media change or the rest of the assay. Organoids in 3-dimensional Matrigel patties were imaged and comprehensively counted while growing using a 2x objective for bright field imaging on a Keyence BZ-X800 microscope. For each well, the Quick Full Focus tool was used to take images across multiple positions on the Z-axis then pick the optimal in-focus position for each pixel, resulting in the full field of organoids appearing in focus. Following 5-FU treatment, live organoids were defined as having a smooth outer barrier visible around the full body of the organoid. Fully opaque black organoids were also excluded as this precluded identifying any living cells. Organoids were counted by hand using the Cell Counter feature on Fiji software. The same four wells were counted for each condition immediately prior to adding 5-FU and again two days following the washout of 5-FU. Data is shown as percentage of organoids surviving after treatment compared to before treatment. Statistical differences were calculated using two tailed Student’s T-tests for each condition.

### Irradiation

Matrigel-plated organoids were cultured in a Corning Costar cell culture plate. Four days following plating, the cells were administered 40 Gy of irradiation (at 2.98 Gy/min) using a Small Animal Radiation Therapy (SmART+) X-Ray Irradiator with no collimator and a 2mm Aluminum filter. Following irradiation, cells were immediately returned to the 37 °C with 5% CO_2_ and imaged daily to monitor organoid survival.

## Abbreviations used in this paper

5-FU: 5-Fluorouracil
CDKN2A: Cyclin-Dependent Kinase Inhibitor 2A
CRC: colorectal cancer
DAPI: 4′,6-diamidino-2-phenylindole
Dox: doxycycline
EdU: 5-Ethynyl-2’-deoxyuridine
Ind: inducible
IR: irradiation
ISC: intestinal stem cell
KO: knockout
LGR5: leucine-rich repeat-containing G protein-coupled receptor 5
SOX9: SRY-box transcription factor 9
rISC: reserve intestinal stem cell
RNAseq: ribonucleic acid sequencing
SI: small intestine
TA: transit amplifying
TGFB: Transforming Growth Factor Beta
WT: Wildtype

## Data Transparency statement

Sequencing data sets are available on the NCBI Gene Expression Omnibus under accession number GSE341884. Materials used within this study will be made available to qualified investigators upon reasonable request.

---

**Supplemental Figure 1: Uncropped Western Blots**

**A)** Uncropped Western Blot for Figure 1E (SOX9, red; Actin, Green). **B)** Uncropped Western Blot for Figure 1G (SOX9, red; Actin, Green).

