## Supplementary figures and images for "SOX9-mediated G1 elongation confers reserve stem cell-associated injury resistance in human intestinal stem cells"

### Supplemental Figure 1

A

Figure 1E

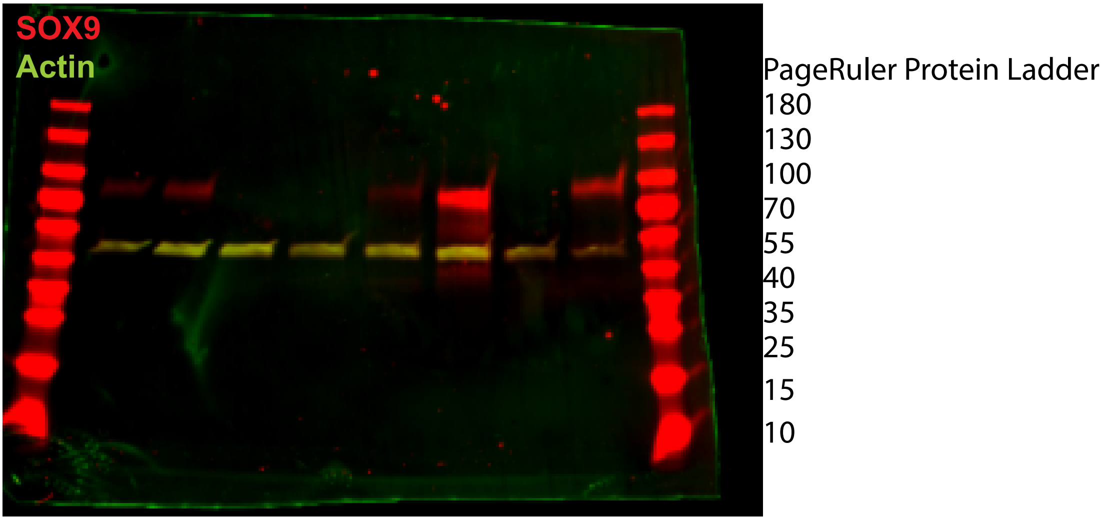


B

Quantified in Figure 1G

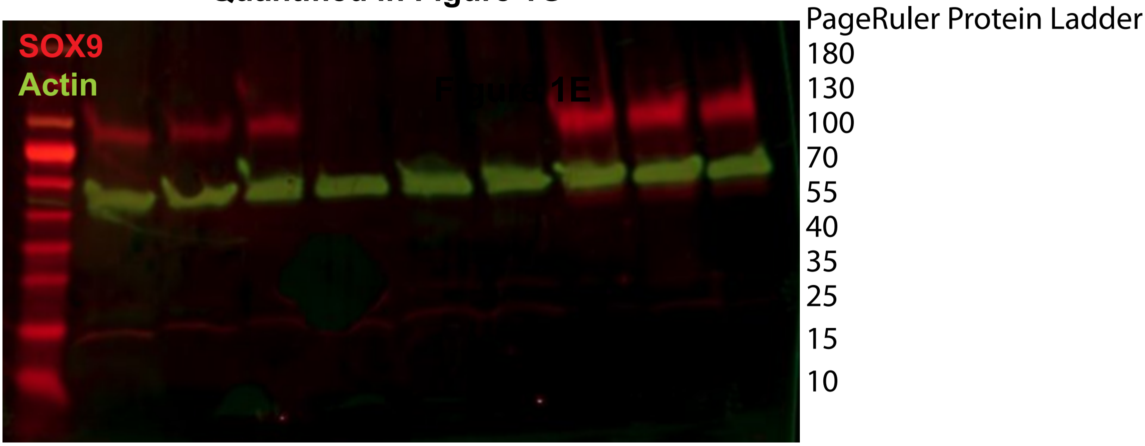
